# Molecular dynamics of the interaction between the ALS/FTD-associated (GGGGCC)n RNA G-quadruplex structure and the three RRM domains of hnRNP H

**DOI:** 10.1101/2023.05.24.541672

**Authors:** Marvin Jericho Cava, Junie B Billones, Josephine Galipon

## Abstract

Hexanucleotide repeat expansions (HRE), located in the first intron of chromosome 9 open reading frame 72 (C9orf72) are the most common genetic abnormality associated with amyotrophic lateral sclerosis (ALS) and frontotemporal dementia (FTD). Presence of the HRE may cause various effects to neuronal cells, leading to pathogenicity. One of these is the sequestration of RNA-binding proteins by three-quartet parallel RNA G-quadruplexes (RG4s) formed from repeated (GGGGCC)n sequences on the sense transcripts of the HRE. Multiple studies imply a major role of the sequestration of heterogeneous nuclear ribonucleoprotein H (hnRNP H) in the pathology of ALS/FTD. In this study, molecular docking and molecular dynamics (MD) were used to simulate the interaction of the three RNA recognition motifs (RRMs) of hnRNP H with the RG4. Molecular Mechanics with Generalised Born and Surface Area Solvation (MM-GBSA) and hydrogen bonding analyses of MD simulations were performed. The MM-GBSA analyses revealed that Arg29, Arg150, and Arg299 are important contributors to the binding, consistent with previous observations of arginine-mediated binding of protein to RNA. In addition, our results point to a previously unknown role of the stretch of residues from Lys72 to Tyr82 on hnRNP H for binding the (GGGGCC)n RG4, forming a hydrogen bonding hotspot. Interestingly, the identified residues are not located in the beta sheet, as would be expected of RRMs in general, suggesting that the binding of hnRNP H to this pathological RG4 may be specifically targeted. This has implications for future *in vitro* studies including but not limited to mutational analysis of these mentioned residues as well as drug development to prevent the sequestration of hnRNP H in ALS/FTD.

## INTRODUCTION

Amyotrophic lateral sclerosis (ALS) and frontotemporal dementia (FTD) are two distinct devastating neurological disorders now believed to lie on a continuous disease spectrum (Conlon et al., 2016; Ling et al., 2013). These disorders can occur alone or together in families, suggesting a genetic link (Conlon et al., 2016; McEachin et al., 2020). In other cases, ALS and FTD may also occur sporadically in individuals (Conlon et al., 2016). ALS is characterized by the degeneration and eventual loss of upper and lower motor neurons (Chou & Norris, 1993). Patients with ALS are characterized by a variety of symptoms including loss of limb function, and difficulty with speech and swallowing (Morgan & Orrell, 2016). Progression of the disorder leads to respiratory failure (Morgan & Orrell, 2016). On the other hand, FTD is characterized by atrophy of the frontal and anterior temporal lobes of the brain, as well as degeneration of the striatum (Snowden et al., 2002). In effect, patients may experience changes in behavior, language presentation, and personality (Neary et al., 2000). In many clinical and pathological phenotypes, the symptoms of ALS and FTD overlap to some extent. For instance, ALS-like motor dysfunction is also manifested in up to 40% of FTD patients, and FTD-like cognitive impairment in up to 50% of ALS patients (Burrell et al., 2011; Thomas et al., 2013).

Hexanucleotide repeat expansions (HRE), (GGGGCC)n located in the first intron of chromosome 9 open reading frame 72 (C9orf72) are the most common genetic abnormality associated with ALS and FTD, accounting for up to 40% of familial ALS/FTD, 5-10% of sporadic ALS, and 4-21% of sporadic FTD (Konopka & Atkin, 2018). A study by DeJesus-Hernandez et al. (2011) determined that C9orf72 in asymptomatic individuals contained 2-23 repeats, while those in ALS and FTD patients contained around 700-1600 repeats. Another paper published side by side in the same journal issue reports 30 repeats as the pathogenicity threshold (Renton et al., 2011). This was confirmed by subsequent *in vitro* studies (Xu et al., 2013; Lee et al. 2013; Haeusler et al. 2014).

Two non-exclusive mechanisms have been proposed as to how the C9orf72 HRE causes disease: loss of function of C9orf72, and gain of toxicity caused by the HRE (Pang & Hu, 2021). The loss of function of C9orf72 in ALS and FTD patients may be attributed to the formation of G-quadruplex (G4) structures and RNA-DNA hybrids known as R-loops, which can lead to transcriptional stalling resulting in truncated HRE-containing abortive transcripts (Haeusler et al., 2014; Y. B. Lee et al., 2013). G4s and R-loops have also been associated with other genetic diseases involving repeat expansions (Konopka & Atkin, 2018). In the current study, we aim to provide more insight on the mechanism of gain of toxicity.

There are a variety of ways through which the C9orf72 HRE may cause toxicity. Firstly, it can be bidirectionally transcribed, hence, producing both sense and antisense transcripts (Zu et al., 2013). Both sense and antisense transcripts have been reported to be translated into dipeptide repeat proteins through a mechanism called repeat-associated non-ATG-initiated translation (Mori et al., 2013; Zu et al., 2013). Both sense and antisense transcripts of the C9orf72 HRE have also been shown to form RNA foci (Conlon et al., 2016; DeJesus-Hernandez et al., 2011; Haeusler et al., 2014; Y. B. Lee et al., 2013; Zu et al., 2013). Aside from forming RNA foci, the sense transcript of the HRE was also determined to preferentially bind some RNA-binding proteins (RBPs) *in vivo* (Haeusler et al., 2014; Y. B. Lee et al., 2013). The HRE was also associated with widespread misregulation of alternative splicing, and alternative polyadenylation in the brains of the patients, further supporting the involvement of RBPs in the pathogenicity of the C9orf72 HRE (Prudencio et al., 2015).

According to Conlon et al. (2016), the RNA-splicing protein hnRNP H (heterogeneous nuclear ribonucleoprotein H) is the major protein bound to the C9orf72 HRE sense transcript. It is hypothesized that the formation of RNA G-quadruplexes (RG4) from the (GGGGCC)n repeats leads to the sequestration of hnRNP H, preventing it from carrying out its function in the splicing of other RNA substrates, and thus, leading to anomalies in alternative splicing (Conlon et al., 2016; Y. B. Lee et al., 2013). Prudencio et al. (2015) reported that multiple RNA-binding motifs affected in alternative splicing changes in the brains of HRE-carrying ALS patients corresponded to known hnRNP H binding sequences. Conlon et al. (2016) further analyzed alternative splicing events by comparing ALS and non-ALS post-mortem cerebellum samples, and reported a decrease in exon inclusion in 11 out of 15 identified hnRNP H splicing targets in ALS. Prior to this, Lee et al. (2013) had shown that the transfection of 72 repeats (GGGGCC)n into SH-SY5Y neuronal precursor cells decreased the inclusion of exon 7 into TARBP2 gene transcripts, and show that a similar effect is obtained by hnRNP H gene knockdown.

All of the above imply a major role of hnRNP H sequestration in the pathology of ALS/FTD. However, the mechanism of interaction between hnRNP H protein and the (GGGGCC)n RG4 structure remains poorly understood. In this study, molecular docking and molecular dynamics (MD) were used to simulate the interaction of hnRNP H with the three-quartet RG4 formed from the (GGGGCC)n repeats. Our results suggest that Arg29, Arg150, Arg299, Phe120, and the stretch of residues from Lys72 to Tyr82 are important contributors to the binding. Interestingly, the identified residues are not located in the beta sheet, as would be expected of RRMs, suggesting that the binding of hnRNP H to this pathological RG4 may be specifically targeted (Cañadillas & Varani, 2003; Deka et al., 2005; Cléry & Allain, 2011). This has implications for future *in vitro* studies including but not limited to mutational analysis of these mentioned residues as well as drug development to prevent the sequestration of hnRNP H in ALS/FTD.

## RESULTS

### MD-based modelling of hnRNP H RRMs

While the hnRNP H protein contains three RNA-recognition motifs (RRMs), only RRM1 and RRM2 have experimentally determined 3D structures publicly available on RCSB PDB [rcsb.org] (Ramelot et al., 2012; Meagher & Stuckey, 2018; Berman, 2000). There are two 3D models of hnRNP H published in the mentioned repository, both of which are for the human variant. One of these models includes RRM1 [PDB ID: 2LXU] only, while the other includes both RRM1 and RRM2 [PDB ID: 6DHS] (Ramelot et al., 2012; Meagher & Stuckey, 2018). To circumvent the lack of an experimentally confirmed 3D model for RRM3, we retrieved the RRM3 structure from AlphaFold-predicted model publicly available at the AlphaFold Protein Structure Database (alphafold.ebi.ac.uk/; Jumper et al., 2021; Varadi et al., 2022). As seen in Figure 1A, apart from the three RRMs, most of the protein is predicted to be relatively unstructured. To confirm the validity of the AlphaFold-predicted RRM3 domain structure, the RRM1 and RRM2 domains were extracted and superimposed with the experimentally validated RRM1 and RRM2 domains, respectively. As shown in Figure 1B-D, the values of the root-mean-square deviation of atomic positions (RMSD) between the AlphaFold prediction and the published crystal structures were sufficiently low for both RRM1 (0.657 Å and 0.804 Å) and RRM2 (0.661 Å), considering that the expected value for homologous proteins is 0.62∼2.31 Å (Chothia & Lesk, 1986). In addition, all three RRMs had high sequence coverage in the AlphaFold database, resulting in high per-residue confidence scores, expressed as the predicted local distance difference (pLDDT) in Figure 1A, suggesting that the prediction for RRM3 is likely to be accurate.

**Figure 1.**
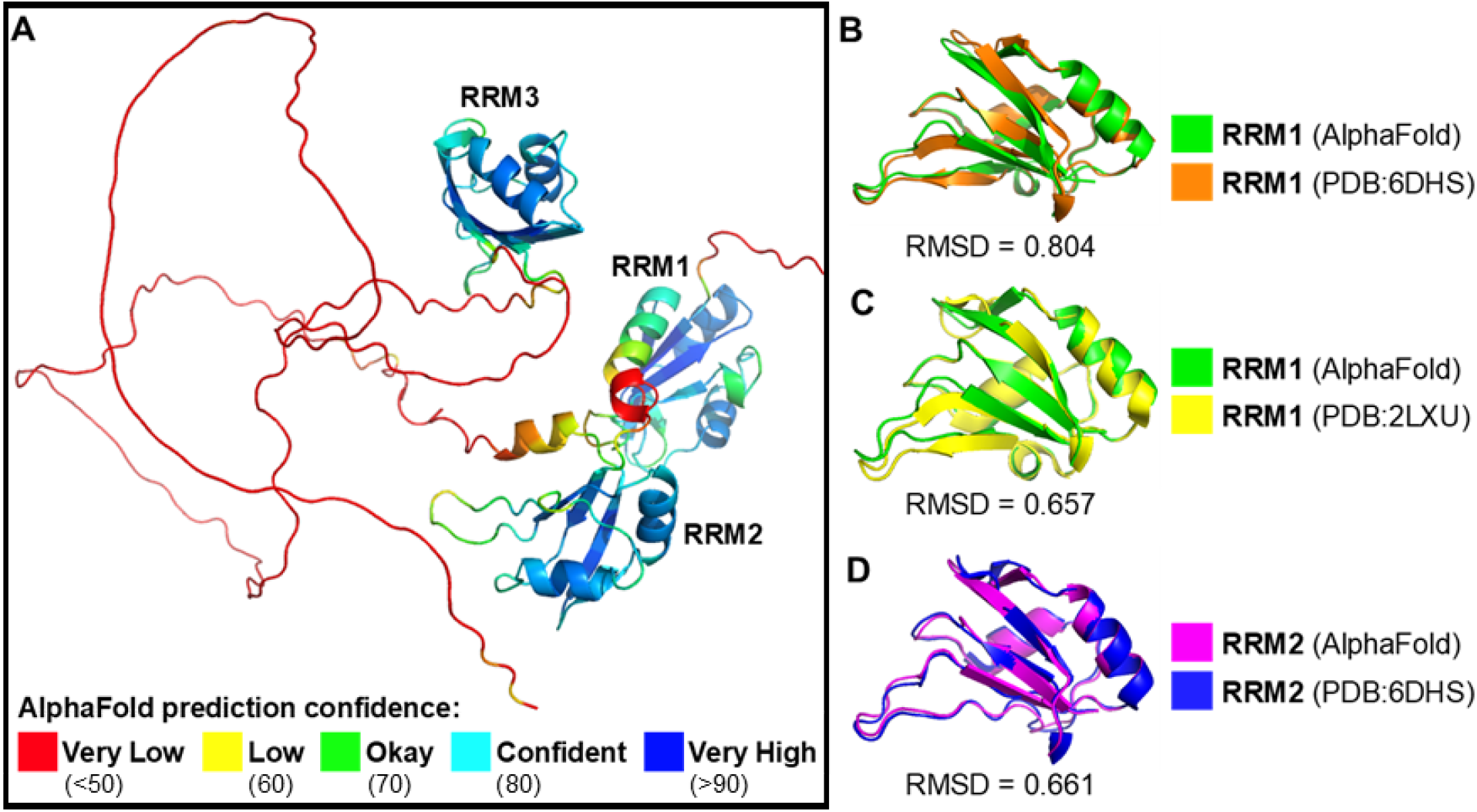
(A) AlphaFold-predicted model of hnRNP H. Red region represents the amino acids leading to RRM3 from the end of RRM2. (B, C, D) Superimposition of experimentally-determined structures of RRM1 and RRM2 onto their respective AlphaFold-predicted structures.

As shown in Figure 1A, the large size of the hnRNP H protein and its long unstructured loops make full-protein simulations difficult. Therefore, to enable molecular dynamics simulations, we initially assumed some degree of interaction between RRM1-2 and RRM3; this was supported by the low predicted aligned error (PAE) for all three RRMs (Figure S1), showing that they are predicted to be positioned reproducibly relative to one another. The RRM3 of hnRNP H was docked onto the published crystallography structure of the connected RRM1-2 (PDB ID: 6DHS) via HDOCK Server (Huang & Zou, 2008; Yan et al., 2020; Yan, Wen, et al., 2017; Yan, Zhang, et al., 2017). We chose HDOCK for its performance in the Seventh Edition of the Critical Assessment of Prediction of Interactions (CAPRI), where it ranked second among ten competing protein-protein docking servers (Lensink et al., 2020). HDOCK is sometimes advantageous over its top-ranked competitor ClusPro, because HDOCK includes both template-based docking and free docking in its algorithm, and the availability of templates may boost the accuracy of HDOCK on certain protein-protein docking tasks (Yan et al., 2020b; Yan, Zhang, et al., 2017). For docking the RRM3 of hnRNP H onto RRM1-2, template-based docking was included.

According to CAPRI’s standards, 60 % of the time, the “true” docking pose is included in HDOCK’s top 10 predictions. Even then, the “true” docking pose may not be of optimum quality. To refine the docking results, we did 50 nanoseconds (ns) molecular dynamics (MD) simulations with the top ten docking poses from HDOCK under a ff19SB (Amber) forcefield. In theory, this step should deconstruct docking poses which are not energetically favorable and optimize “true” docking poses into a more energetically favorable state. After the 50 ns MD simulations, we did an additional 0.1 ns MD simulation starting from the final snapshots of the 50 ns simulations. The purpose of these simulations was to sample minute movements of the structure (rather than obtaining an energetically favorable conformation) for subsequent Molecular Mechanics with Generalised Born and Surface Area Solvation (MM-GBSA) analysis.

It must also be noted that there is no certainty as to how the RRM3 interacts with RRMs 1 and 2 in a cellular environment, and that all poses predicted by HDOCK may be possible solutions *in vivo*. Re-ranking via MM-GBSA was done as an attempt to gauge which configurations would be the most likely from the perspective of thermodynamics. Table 1 shows the results of the HDOCK and MM-GBSA analyses.

**Table 1.** Results of the MM-GBSA analysis. All units are reported in kcal/mol. Docking poses are illustrated in Figure S2.

| **HDock Rank** | $\Delta E_{elec}$ | $\Delta E_{vdw}$ | $\Delta G_{GB}$ | $\Delta G_{SA}$ | **Total** | **MM-GBSA Rank** |
| --- | --- | --- | --- | --- | --- | --- |
| 1 | -385.30 | -89.75 | 403.82 | -14.46 | -85.69 | 1 |
| 2 | -124.26 | -22.41 | 136.61 | -3.81 | -13.88 | 10 |
| 3 | -75.27 | -36.64 | 101.42 | -5.28 | -15.26 | 9 |
| 4 | -210.11 | -59.44 | 245.04 | -8.33 | -32.84 | 8 |
| 5 | -377.73 | -69.19 | 398.77 | -10.88 | -59.03 | 2 |
| 6 | -212.50 | -75.07 | 248.48 | -11.26 | -50.36 | 6 |
| 7 | -344.43 | -73.96 | 372.12 | -11.71 | -57.97 | 3 |
| 8 | -280.61 | -76.24 | 313.16 | -11.87 | -55.56 | 5 |
| 9 | -350.00 | -57.36 | 375.79 | -8.43 | -40.80 | 7 |
| 10 | -372.82 | -69.73 | 396.63 | -10.79 | -56.12 | 4 |

The docking pose which ranked first in both scoring functions (HDOCK and MM-GBSA) is predicted to have the most unfavorable ΔG_GB_, but was also predicted to have the most favorable ΔE_elec_, ΔE_vdw_, and ΔG_SA_. Adding the ΔG_GB_ and ΔG_SA_ together results in a positive value, suggesting an unfavorable reorganization of solvent particles when the RRMs of hnRNP H are oriented as in HDOCK Rank 1. This is however greatly compensated by the highly negative ΔE_elec_ and ΔE_vdw_, which represent the favorable interaction of the RRM3 with RRM1 and RRM2. Overall, HDOCK Rank 1 was predicted to be the most favorable among the ten docking poses, even when the standard deviation among the MD snapshots in the trajectories were considered–see Figure 2. Moreover, the stability of HDOCK Rank 1 can be inferred from the RMSD and RG (radius of gyration) plots of the 50 ns MD trajectory–see Figures S3 and S4 (same may be true for some of the other poses).

**Figure 2.**
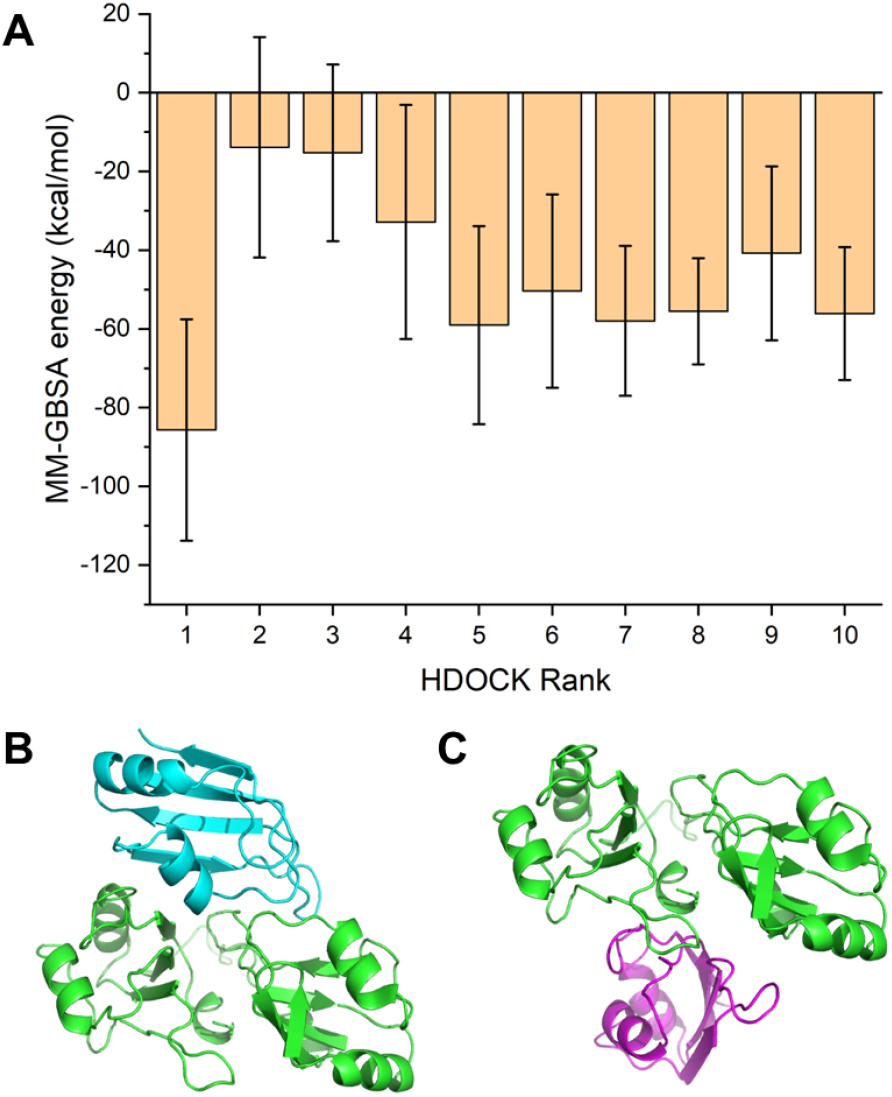
(A) MM-GBSA energies of the ten docking poses. The error bars represent the standard deviation among 1000 frames spread across 0.1 ns of MD simulation. (B) HDOCK Rank 1 docking pose (Model A). RRM3 is colored blue. (C) HDOCK Rank 5 docking pose (Model B). RRM3 is colored purple.

While there is no discrepancy in the topmost-ranked poses for both HDOCK and MM-GBSA, HDOCK Rank 5 ranked second in the MM-GBSA re-ranking. It can be observed from the RMSD and RG plots (Figures S3 and S4, respectively) that there was movement throughout its (HDOCK Rank 5) MD trajectory, but not to the point of deconstruction (as is the case with the HDOCK Rank 3 docking pose–see Figures S5 and S6). As in HDOCK Rank 1, adding the ΔG_GB_ and ΔG_SA_ of HDOCK Rank 5 results in a positive value, but this was greatly compensated by the highly negative ΔE_elec_ and ΔE_vdw_.

Docking poses corresponding to HDOCK Ranks 1 and 5, being the two most favorable among the predicted docking poses (based on MM-GBSA) were the models used to predict the interactions of the RG4 with hnRNP H. The docking models obtained from HDOCK were the ones used for subsequent procedures. From here onwards, the chosen docking poses shall be referred to as Model A (for HDOCK Rank 1) and Model B (for HDOCK Rank 5), respectively.

### MD-based modelling of C9orf72 HRE RNA G-quadruplex

To our knowledge, the 3D structure of (GGGGCC)n RG4 has not yet been experimentally determined. Thus, in the current study, we predicted the RG4 3D structure from the structure of another RG4-forming sequence via an MD-based homology modelling approach. Other studies have predicted the 3D structure of other RG4s using a similar method (Carvalho et al., 2020; Mulholland et al., 2020; Santos et al., 2021). In contrast to the other studies which used additive forcefield for MD simulations of G4s, in this study, we opted for a polarizable forcefield, Drude, which is known for its better performance in the MD simulation of G4s (Li et al., 2021).

Haeusler et al. (2014) determined that the C9orf72 HRE RNA preferentially adopts a three-quartet parallel G4 topology (Figure 3A) at physiological conditions. Four repeats of the HRE RNA subjected to circular dichroism spectroscopy showed the signature spectra for parallel G4s while RNase protection assay provided evidence for a three-stacked parallel-stranded G4 topology (Haeusler et al., 2014). Thus, as a basis for an MD-based homology model, we used the experimentally determined structure of the parallel three-quartet intramolecular G4 formed from human telomeric DNA (PDB ID: 1KF1; Parkinson et al., 2002). Adenine bases in the model were changed into guanine and thymine bases were changed into cytosine (Figure 3D). An -OH groupement was then added to each deoxyribose in the DNA backbone to convert the structure to RNA. The model then underwent 300 ns MD simulation under a Drude polarizable forcefield (Lamoureux et al., 2006; H. Yu et al., 2010; W. Yu et al., 2013).

**Figure 3.**
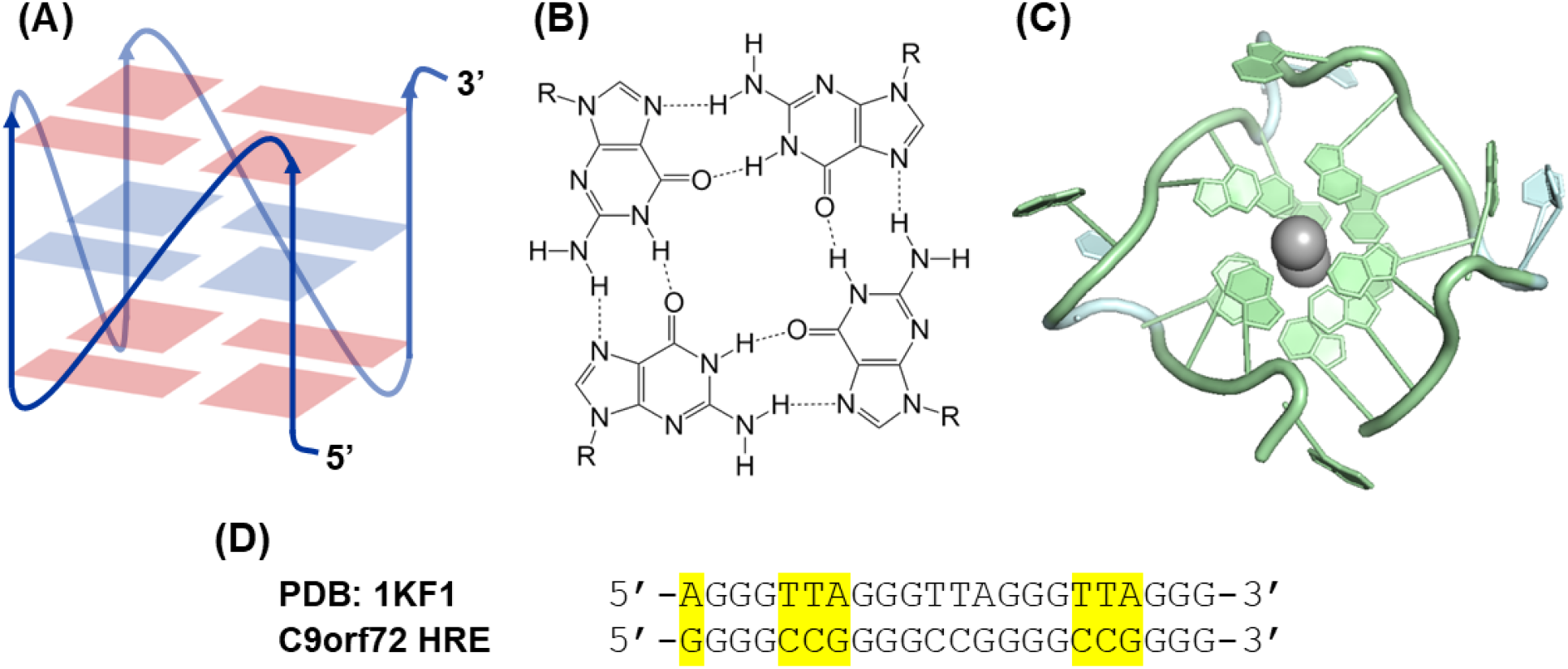
(A) Diagram of a three-quartet intramolecular parallel G4. (B) Guanine quartet. (C) Visualization of the C9orf72 HRE RG4 model produced in this study. (D) Nucleotide sequences of the template model compared to the C9orf72 HRE.

Judging from the RMSD and RG plots of the MD trajectory (Figure S7), the structure of the RG4 stabilized at around 100 ns of MD simulation. Moreover, from the RMSF plot for the last 100 ns of the MD simulation (Figure S8), the guanine residues from the RG4 quartets had low RMSF, which means that the quartets are stable. Two K^+^ ions were also stably positioned in the ion channel of the RG4 during the same time frame (Figure S9). The final snapshot of the 300 ns MD simulation was extracted for use as a model for docking of the C9orf72 RG4 onto the RRMs of hnRNP H.

### Docking of the RG4 onto hnRNP H and MD simulation of docking poses

The predicted RG4 model was docked using HDOCK onto two different models of hnRNP H (Models A and B). We selected HDOCK for its good performance in three different protein-RNA docking benchmarks: Pérez-Cano et al. (2012), Nithin et al. (2017), and Huang & Zou (2013) benchmarks. When the top prediction is considered, HDOCK’s success rates in the three benchmarks were 33.3%, 33.3% and 52.0%, respectively (Yan, Zhang, et al., 2017). When the top 10 predictions are considered, the success rates increased to 54.5%, 51.5% and 64.0%, respectively (Yan, Zhang, et al., 2017).

The top 10 docking results for each of the two models of hnRNP H were subjected to 170 ns MD simulations under Drude polarizable forcefield. This is to optimize the docking poses as was previously done in the hnRNP H modelling. Since the MD simulations in this section of our study involve an RG4, we used a Drude polarizable forcefield for the same reason discussed in the previous section.

As previously mentioned, the models obtained from HDOCK (Ranks 1 and 5, hereby referred to as Models A and B, respectively) were used in this section rather than the final snapshots of the previous MD simulations of the docking poses. The rationale for making this choice was that even though the structures are likely to stabilize slightly differently in the Drude polarizable forcefield, the initial pose and the stabilized final snapshots are not significantly different (RMSD = 0.36 Å for Model A, and 0.53 Å for Model B, respectively). Docking of the RG4 into the hnRNP H RRMs was also expected to affect how the RRMs would reorient towards more energetically favorable conformations in an MD simulation. The MD simulations which were performed for this section of the current study are sufficiently long (170 ns), so that all components of the simulated systems had time to stabilize. Indeed, based on the RMSD and RG plots, almost all the trajectories for the top ten docking poses of the RG4 onto hnRNP H Models A and B were stable at least by the last 30 ns of the MD simulations (Figures S10-S13).

Subsequent to these 170 ns MD simulations, we did 1 ns MD simulations starting from the final snapshots of the previous 170 ns simulations under ff19SB (for the protein receptor) and OL3 (for the RNA ligand) Amber forcefields. The purpose of these simulations was to sample minute movements of the structure (rather than obtaining an energetically favorable conformation) for subsequent MM-GBSA analysis. As described above, MM-GBSA was also performed here to re-rank the docking poses. Table 2 shows the results of the MM-GBSA analysis.

**Table 2.** Results of the MM-GBSA analysis. All units are reported in kcal/mol. Docking poses are illustrated in Figures 4 and 5.

| hnRNP H Model | HDOCK Rank | $\Delta E_{elec}$ | $\Delta E_{vdw}$ | $\Delta G_{GB}$ | $\Delta G_{SA}$ | Total | MM-GBSA Rank (Global) |
| --- | --- | --- | --- | --- | --- | --- | --- |
| Model A | 1 | 98.37 | -79.11 | -93.68 | -8.85 | -83.27 | 7 |
| Model A | 2 | 53.91 | -58.45 | -53.26 | -7.52 | -65.32 | 12 |
| Model A | 3 | 31.91 | -74.60 | -28.12 | -8.30 | -79.11 | 8 |
| Model A | 4 | -65.15 | -55.12 | 63.76 | -5.69 | -62.20 | 14 |
| Model A | 5 | 86.05 | -61.06 | -82.85 | -7.97 | -65.83 | 11 |
| Model A | 6 | -59.59 | -44.51 | 57.97 | -5.96 | -52.09 | 17 |
| Model A | 7 | -9.51 | -72.16 | 12.00 | -8.78 | -78.45 | 9 |
| Model A | 8 | 40.55 | -76.44 | -38.5 | -10.46 | -84.85 | 6 |
| Model A | 9 | 14.03 | -62.57 | -13.42 | -7.58 | -69.54 | 10 |
| Model A | 10 | -40.14 | -57.32 | 40.64 | -5.93 | -62.75 | 13 |
| Model B | 1 | 68.99 | -15.43 | -68.17 | -2.42 | -17.03 | 20 |
| Model B | 2 | 25.61 | -80.87 | -23.13 | -9.58 | -87.97 | 4 |
| Model B | 3 | 39.15 | -94.40 | -34.18 | -11.32 | -100.75 | 2 |
| Model B | 4 | 9.29 | -82.19 | -7.72 | -9.20 | -89.82 | 3 |
| Model B | 5 | 17.62 | -50.48 | -16.69 | -6.00 | -55.55 | 16 |
| Model B | 6 | 21.35 | -77.41 | -19.79 | -9.50 | -85.35 | 5 |
| Model B | 7 | 33.6 | -48.12 | -31.86 | -5.35 | -51.73 | 18 |
| Model B | 8 | 100.54 | -47.27 | -95.3 | -5.52 | -47.55 | 19 |
| Model B | 9 | 48.93 | -56.42 | -47.38 | -6.40 | -61.27 | 15 |
| Model B | 10 | 9.76 | -95.72 | -5.71 | -10.04 | -101.71 | 1 |

All the ΔE_vdw_ values in Table 2 are highly negative, indicating that the van der Waals interactions between the hnRNP H and the RG4 are predicted to be favorable. The ΔG_SA_ values are also all negative indicating that the nonpolar component of the energy for the reorganization of solvent particles (around the hnRNP H and the RG4) brought about by the complex-formation is predicted to be favorable. ΔE_elec_ and ΔG_GB_ values, while mostly positive and negative, respectively, are heterogeneous. ΔE_elec_ represents the polar component of the free energy change resulting from electrostatic interactions of the hnRNP H and the RG4. ΔG_GB_ represents the component of the free energy change resulting from electrostatic interactions between the solvent, and the hnRNP H and RG4. Overall, the 20 docking poses are predicted to be favorable as signified by the negative values of the total MM-GBSA energies. This implies that the hnRNP H-RG4 complex may possibly form in vitro and more importantly, in vivo, agreeing with earlier mentioned studies (see Introduction). From Table 2, it can be observed that the first through fifth in the ranking (global) are all docking poses of the RG4 onto Model B, and the sixth through tenth in the ranking are all docking poses of the RG4 onto Model A.

### Identification of amino-acids involved in binding of hnRNP H to the RG4

Figure 6 shows the plots for the per-residue decomposition of the MM-GBSA energies of the top 10 docking poses (top 10 as indicated in the MM-GBSA re-ranking–see Table 2). While per-residue decomposition values were also assigned for the residues of RG4, for the sake of this discussion, the plots show only the simulated residues of hnRNP H. Interestingly, the plots show that all Ranks 1 through 4, together with Rank 8 and Rank 10, have peaks for the range of residues from Lys72 to Tyr82, for Phe120, for Arg150, and for Arg299. It must be noted that the RRM3 has a different orientation in the hnRNP H model used for docking Ranks 1 through 4 than that used for docking Rank 8 and Rank 10. Given that the residues (or stretch of residues) are important to the MM-GBSA energies of the mentioned docking poses, it is reasonable to expect that these residues could be important for the binding of hnRNP H with the RG4 in living cells. Another amino acid of interest is Arg29, for which the plots of Ranks 1 - 5 and Rank 9 also show a peak.

**Figure 4.**
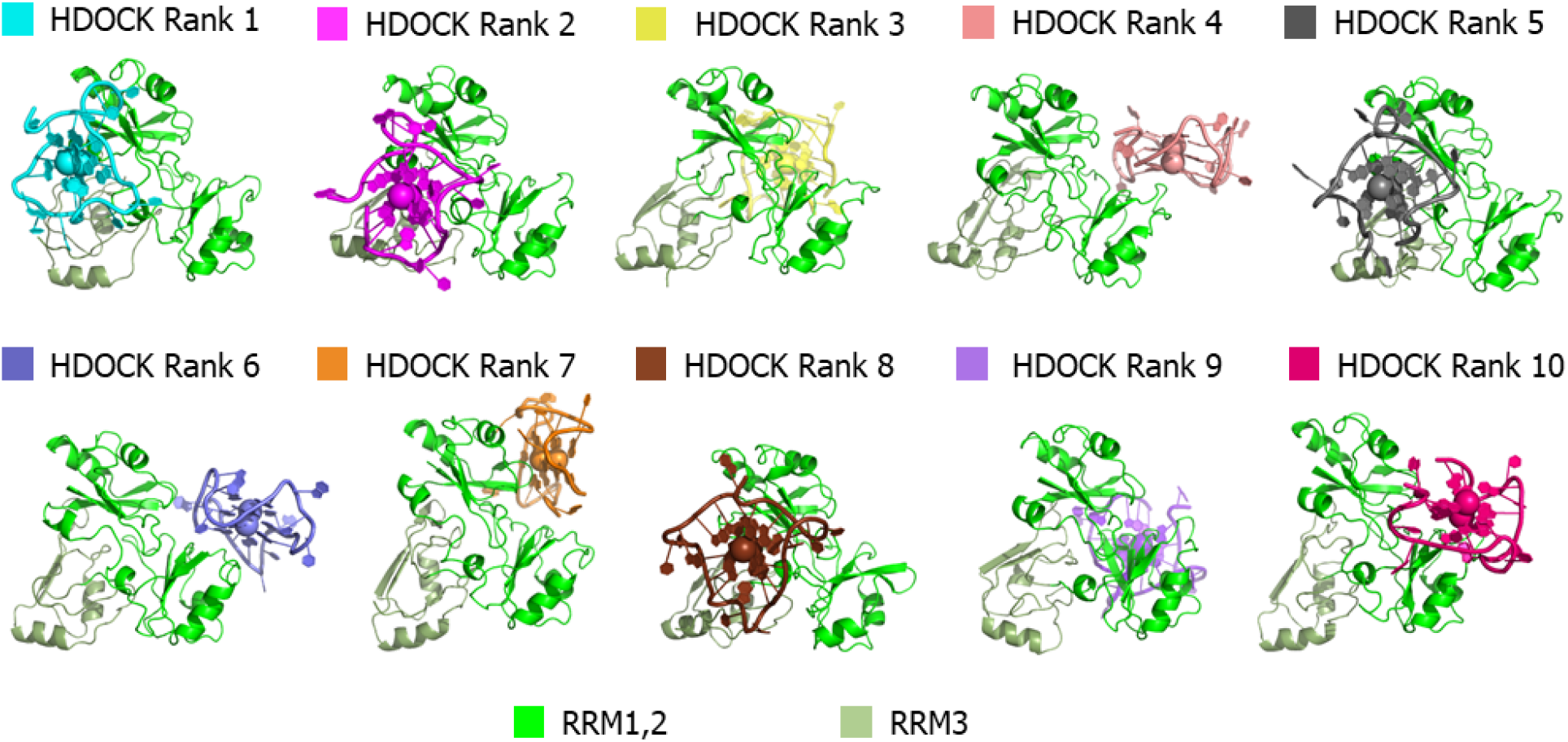
Final snapshots of the 170 ns MD simulations of the docking poses of the C9orf72 RG4 onto hnRNP H Model A.

**Figure 5.**
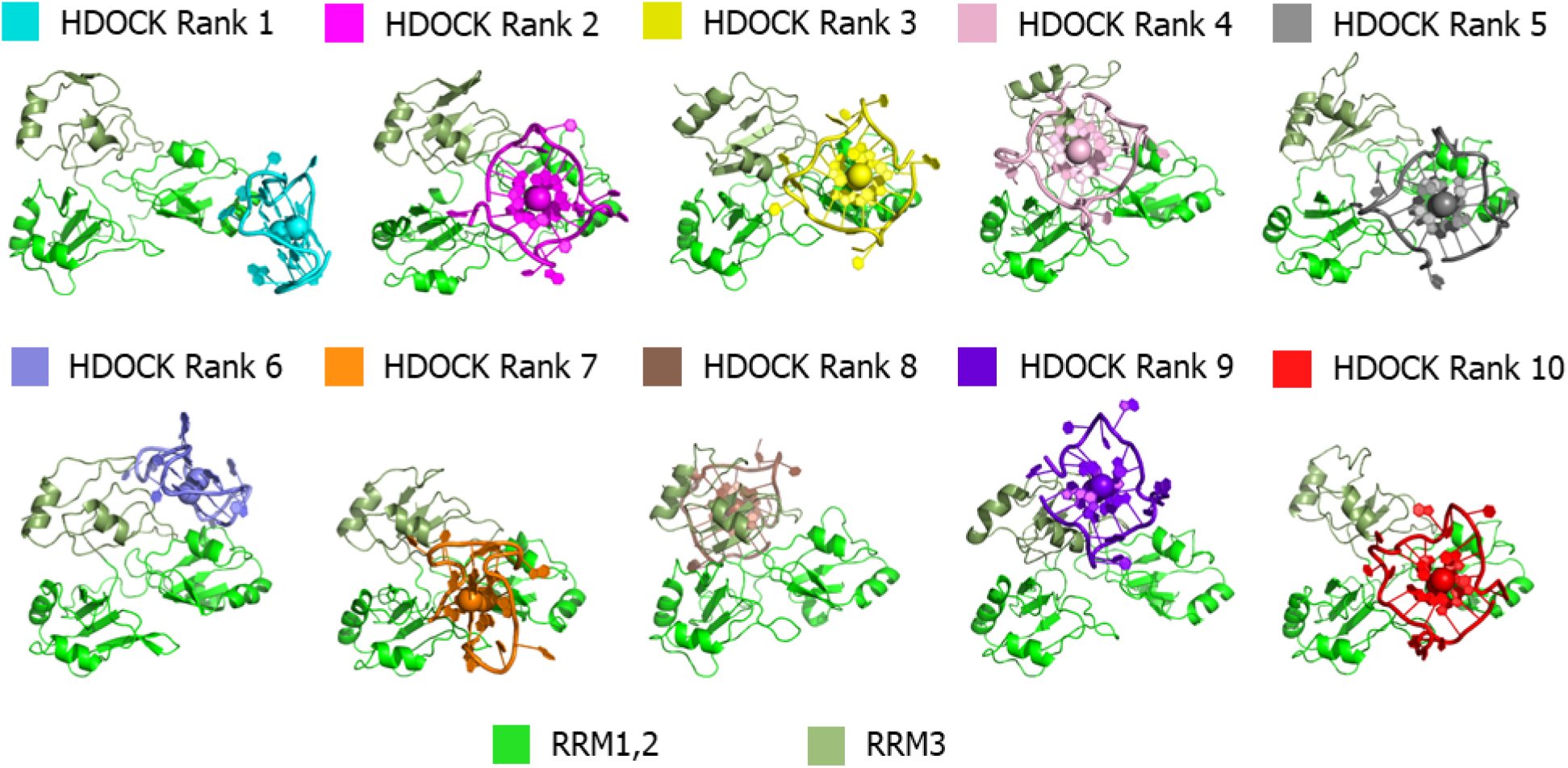
Final snapshots of the 170 ns MD simulations of the docking poses of the C9orf72 RG4 onto hnRNP H model B.

**Figure 6.**
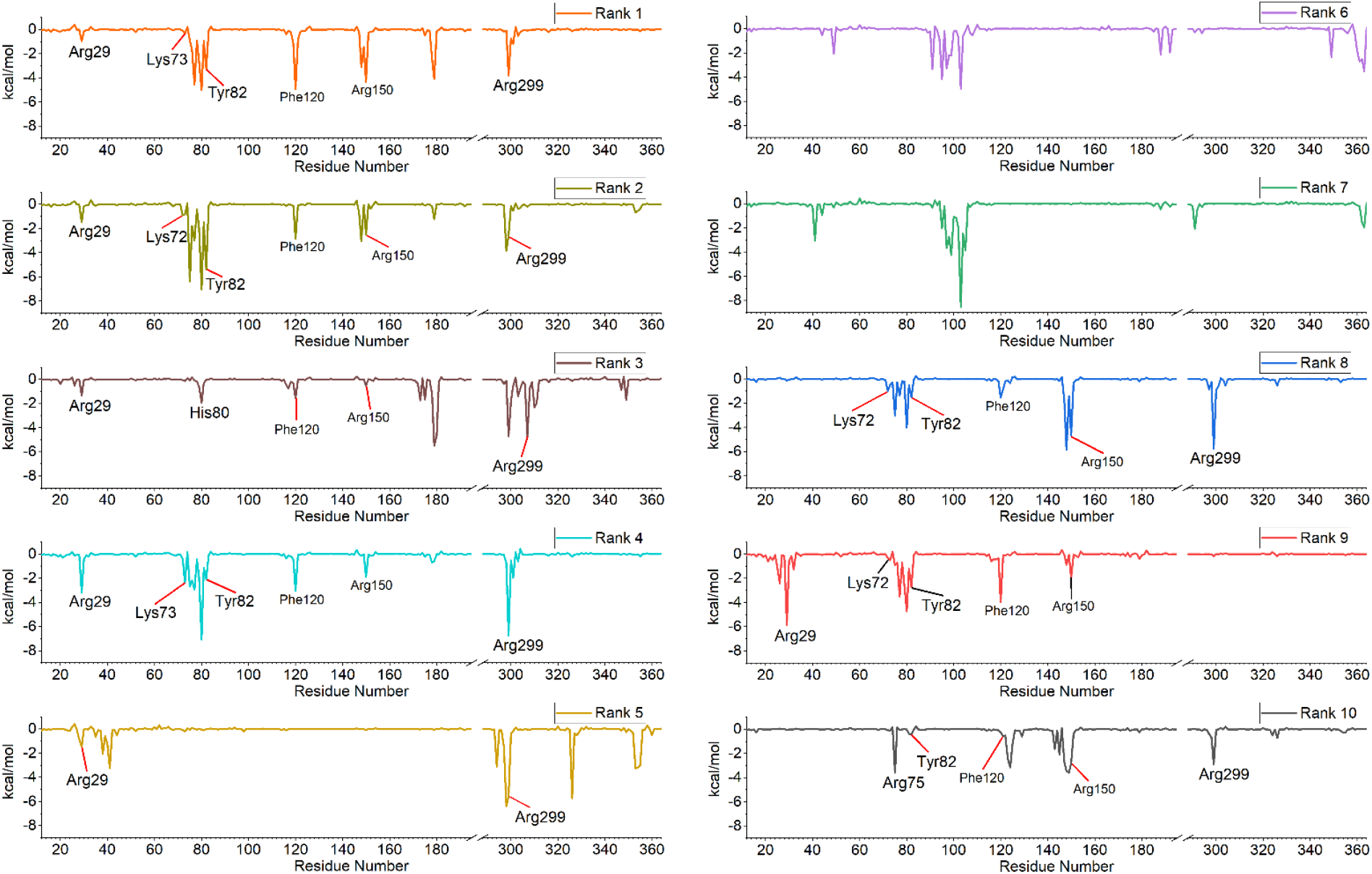
Contributions of the residues of hnRNP H (Model A for Ranks 1-5; Model B for Ranks 6-10) to the MM-GBSA energies of the docking poses.

Notably, three arginine residues, one for each of the RRMs of hnRNP H, are predicted here to be important contributors to the binding of hnRNP H to the RG4 based on the MM-GBSA energies decomposition as illustrated in Figure 6. This finding is consistent with previous findings on the role of arginine residues in the mediation of RNA recognition (Draper, 1999; Chavali et al., 2020). This is thought to be due to their positive charge at physiological conditions, which promotes electrostatic interactions with the negatively charged phosphate backbone of RNA, an interaction that is also dependent on the structural context (Draper, 1999; Chavali et al., 2020). Such “architectural specificity” is likely the case with the three arginine residues described here as indicated by their contribution to the MM-GBSA energies. Other arginine residues (Arg44, Arg49, Arg75, Arg179, Arg188, Arg192, and Arg326) were also predicted in our 50 ns MD simulations of some of the docking poses to form hydrogen bonds with residues of the RG4 (Table S1-10, Figure S15). Mutational analysis is needed to confirm the role of these arginine residues in RNA binding. If mutating an arginine residue to a lysine does not significantly reduce RNA binding, the arginine’s role is simply to contribute a positive charge (Tan & Frankel, 1994).

As for Phe120, it has an aromatic amino acid side chain which can pi-stack with nucleotides.The phenylalanine side chain has no functional groups capable of hydrogen bonding. Only the amine group and the carbonyl group at the main chain can facilitate hydrogen bonding. Thus, the high MM-GBSA predicted to be contributed by Phe120 in the MM-GBSA analyses are likely from factors other than hydrogen bonding. It may be that the conformation of the phenylalanine amino acid residue (adjacent amino acids may be a factor for this) and/or its pi-stacking with nucleotides of the RG4 are energetically favorable.

Hydrogen bonding analysis of the Drude trajectories of the top 10 (global) docking poses also revealed that the stretch of residues from Lys72 to Tyr82, which were previously identified as contributory to the MM-GBSA energies are also predicted to be a hydrogen bonding hotspot–see Figure S15. It may be strategic to focus on this region in future *in vitro* studies probing into the interaction of hnRNP H with the C9orf72 HRE RG4.

Finally, it is worth mentioning that the residues identified here as thermodynamically important in the binding of hnRNP H to the (GGGGCC)n RG4 are distributed across all three RRMs, suggesting that all three RRMs cooperate to bind the RG4. Also, all of them except for Arg29 are in the loops rather than the β-sheets or α-helices. Arg29 is at the α1 region of RRM1; the stretch of residues at Lys72 to Tyr82 at the loop from β1 to α1 of RRM2; Phe 120 at the loop from β1 to α1 of RRM2; Arg150 at the loop from α1 to β2 of RRM2; and Arg299 at the loop from β1 to α1 of RRM3 (Figure S14). These results diverge from observations in classical RRMs where β-sheets serve as primary binding surfaces (Cañadillas & Varani, 2003; Deka et al., 2005; Cléry & Allain, 2011). An *in vitro* study by Dominguez and Allain (2006) also reports interactions involving amino-acids outside of the β-sheets with the two RRMs of hnRNP F and a different G-rich RNA.

## CONCLUSION

We have modeled and studied the interaction of hnRNP H with the C9orf72 HRE RG4 via a series of molecular docking and molecular dynamics experiments. We first predicted how the three RRMs of hnRNP H would be positioned together by docking the RRM3 into the two other RRMs. The docking poses were optimized via molecular dynamics and ranked via MM-GBSA analyses. The top two poses were then used as models for docking the RG4. Docking poses involving both of the top two models were optimized via molecular dynamics, and studied via MM-GBSA analyses and hydrogen bonding analyses. Based on the MM-GBSA analyses, Arg29, Arg150, Arg299, Phe120, and the stretch of residues from Lys72 to Tyr82 were identified as important to the binding of the hnRNP H to the RG4. Results for the hydrogen bonding analyses further show that Lys72 to Tyr82 is a binding hotspot. We suggest that future *in vitro* studies including but not limited to mutational analyses probe these mentioned residues.

## METHODS

### Modelling of hnRNP H RRMs

A model for RRM3 was obtained by splicing residues 289-364 from the AlphaFold-predicted model publicly available at the AlphaFold Protein Structure Database (AlphaFold DB version 2022-06-01, model created with the AlphaFold Monomer v2.0 pipeline; UniProt: P31943) of hnRNP H (Jumper et al., 2021; Varadi et al., 2022). Splicing of the model was performed via PyMOL 2.5. The RRM3 of hnRNP H was then docked into RRMs 1 and 2 via HDOCK Server (Huang & Zou, 2008; Yan et al., 2020a; Yan, Wen, et al., 2017; Yan, Zhang, et al., 2017). Missing residues in the PDB model for RRMs 1 and 2 (PDB ID: 6DHS) were added using PDBFixer version 1.6 (Eastman et al., 2013; Penumutchu et al., 2018). 50 ns MD simulations of each of the top 10 docking poses from HDOCK Server were performed using ff19SB (Amber) forcefield (Tian et al., 2020) via GROMACS 2021.3 (Abraham et al., 2015). Files used for the simulations were prepared using CHARMM-GUI Solution Builder (Jo et al., 2008; J. Lee et al., 2016, 2020). Other than the setting for which simulation software and forcefield were to be used for the MD simulations, default settings for CHARMM-GUI Solution Builder were maintained. The simulation parameters file step5_production.mdp was then edited so that the nsteps parameter is set to 25000000. In preparation for MM-GBSA analyses, we did additional 0.1 ns MD simulations (production only) starting from the final snapshots of the 50 ns simulations. The same input files were used, except that the simulation parameters file step5_production.mdp was edited again, so that the following parameters have these respective values: dt = 0.001; nsteps = 100000; nstxtcout = 100. MM-GBSA analyses were performed via gmx_MMPBSA (v. 1.5.6) to re-rank the docking poses (Valdés-Tresanco et al., 2021; Miller et al., 2012). The following general namelist variable parameters were set for running gmx_MMPBSA: startframe = 1; endframe = 1000; interval = 1; forcefields = “leaprc.protein.ff19SB”; PBRadii = 3; temperature = 303.15. The following Generalized-Born namelist variable parameters were set for running gmx_MMPBSA: igb = 2; extdiel = 80.0; saltcon = 0.15. The top 2 ranking docking poses (MM-GBSA ranking) were later used for docking the RG4 model.

### MD-based modelling of C9orf72 HRE RNA G-quadruplex

As a base for an MD-based homology model, we used the experimentally determined structure of the parallel three-quartet intramolecular G-quadruplex formed from human telomeric DNA (PDB:1KF1; Parkinson et al., 2002). Adenine bases in the model were changed into guanine and thymine bases were changed into cytosine. A 2’ oxygen atom was then added to each deoxyribose in the DNA backbone to convert the structure to RNA. These changes were made in the PDB file via UCSF ChimeraX 1.2.5 (Pettersen et al., 2021). The model then underwent 300 ns MD simulation under Drude polarizable forcefield via OpenMM 7.4.2 (Eastman et al., 2017; Lamoureux et al., 2006; H. Yu et al., 2010; W. Yu et al., 2013). Before the MD simulation under Drude polarizable forcefield, a pre-equilibration step was performed using CHARMM36 additive forcefield via OpenMM (Lee et al., 2016). Input files for the pre-equilibration were generated from the modified PDB file using CHARMM-GUI Solution Builder. Default settings were used. For the MD simulation under Drude polarizable forcefield, input files were generated from the output in the pre-equilibration step using CHARMM-GUI Drude Prepper (Kognole et al., 2022; J. Lee et al., 2016). In the Drude Prepper interface, the PSF file was uploaded as X-PLOR format. The “Setup PBC” option was unticked. System Type was set to “Solution.” To set the length of the production simulation, the variable “nstep” was set to 300000000 by editing the step5_production.inp file. The final snapshot of the 300 ns MD simulation was extracted as PDB file for use as model for docking of the C9orf72 RG4 onto the RRMs of hnRNP H. Imaginary particles incorporated for MD simulation with the Drude forcefield were removed from the final snapshot before proceeding to the docking. Solvent and ions other than those in the ion channel of the RG4 were also removed.

### Docking of the RG4 onto hnRNP H RRMs

For this section, the previously produced models for hnRNP H were used as the receptor molecule, while the previously produced MD-based model of the RG4 was used as the ligand molecule. HDOCK Server was used to dock the RG4 onto the hnRNP H models. Default settings for HDOCK were all maintained.

### Molecular Dynamics

The top 10 docking poses given by the HDOCK server for each of the two hnRNP H models underwent 170 ns MD simulations under Drude polarizable forcefield via OpenMM. Before the MD simulations under Drude polarizable forcefield, a pre-equilibration step was performed using CHARMM36 additive forcefield via OpenMM. Input files for the pre-equilibration were generated using CHARMM-GUI Solution Builder. Default settings were used. For the MD simulation under Drude polarizable forcefield, input files were generated from the output in the pre-equilibration step using CHARMM-GUI Drude Prepper. In the Drude Prepper interface, the PSF file was uploaded as X-PLOR format. The “Setup PBC” option was unticked. System Type was set to “Solution.” To set the length of the production simulation, the variable “nstep” was set to 170000000 in the step5_production.inp file. The final snapshot of the 170 ns MD simulation was extracted as PD file for use in the next procedure. Imaginary particles incorporated for MD simulation with the Drude forcefield were removed. Solvent and ions other than those in the ion channel of the RG4 were also removed.

### Reranking of the Docking Poses

To rerank the docking poses, MM-GBSA analysis of the MD simulated docking poses was performed via gmx_MMPBSA. Before proceeding to the MM-GBSA analysis, the final snapshots of the Drude simulations of the docking poses underwent 1 ns MD simulations under gmx_MMPBSA-compatible Amber forcefields, ff19SB (for the protein receptor) and OL3 (for the RNA ligand) via GROMACS. Input files for the simulations were generated via CHARMM-GUI Solution Builder. Other than the setting for which simulation software and forcefields were to be used for the MD simulations, default settings were maintained. The simulation parameters file step5_production.mdp was edited so that the following parameters have these respective values: nstxtcout = 5000; nstvout = 5000; nstfout = 5000; constraint_algorithm = SHAKE. After the 1 ns MD simulations, MM-GBSA analysis was performed on the produced trajectories. The following general namelist variable parameters were set for running gmx_MMPBSA: startframe = 1; endframe = 100; forcefields = “leaprc.protein.ff19SB,leaprc.RNA.OL3”; temperature = 303.15. The following Generalized-Born namelist variable parameters were set: igb = 7; intdiel = 8.0; saltcon = 0.15.

### Hydrogen bonding analysis

Hydrogen bonding analysis was done in VMD 1.9.4a53 (Humphrey et al., 1996). Selection 1 was set to “protein,” and Selection 2 was set to “nucleic.” Donor-acceptor distance was set to 3.5 Å, and Angle cutoff was set to 30°. “Update selections every frame?” and “Only polar atoms (N, O, S, F)?” options were ticked. Detailed info for “Residue pairs” were calculated. Only the last 30 ns of the 170 ns Drude forcefield MD simulations were used for the hydrogen bonding analysis.

