## Supplementary Data for "Molecular dynamics of the interaction between the ALS/FTD-associated (GGGGCC)n RNA G-quadruplex structure and the three RRM domains of hnRNP H"

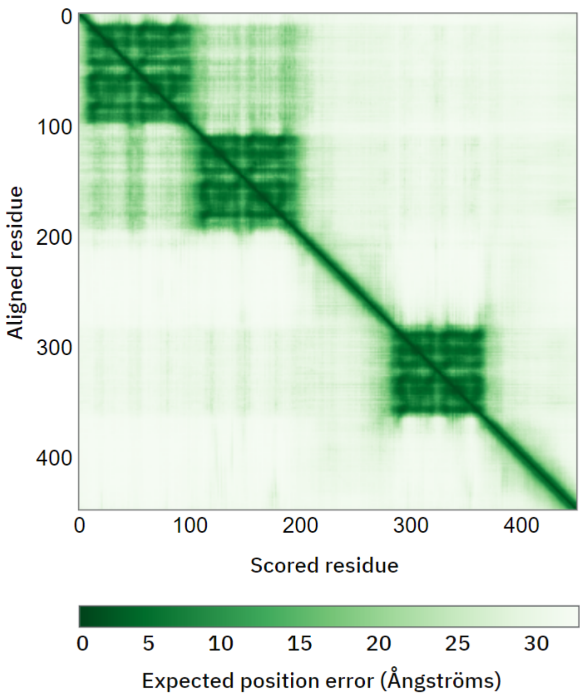


**Figure S1.** Color at position (x, y) indicates AlphaFold's expected position error at residue x, when the predicted and "true" structures are aligned on residue y. Plot from AlphaFold Protein Structure Database.

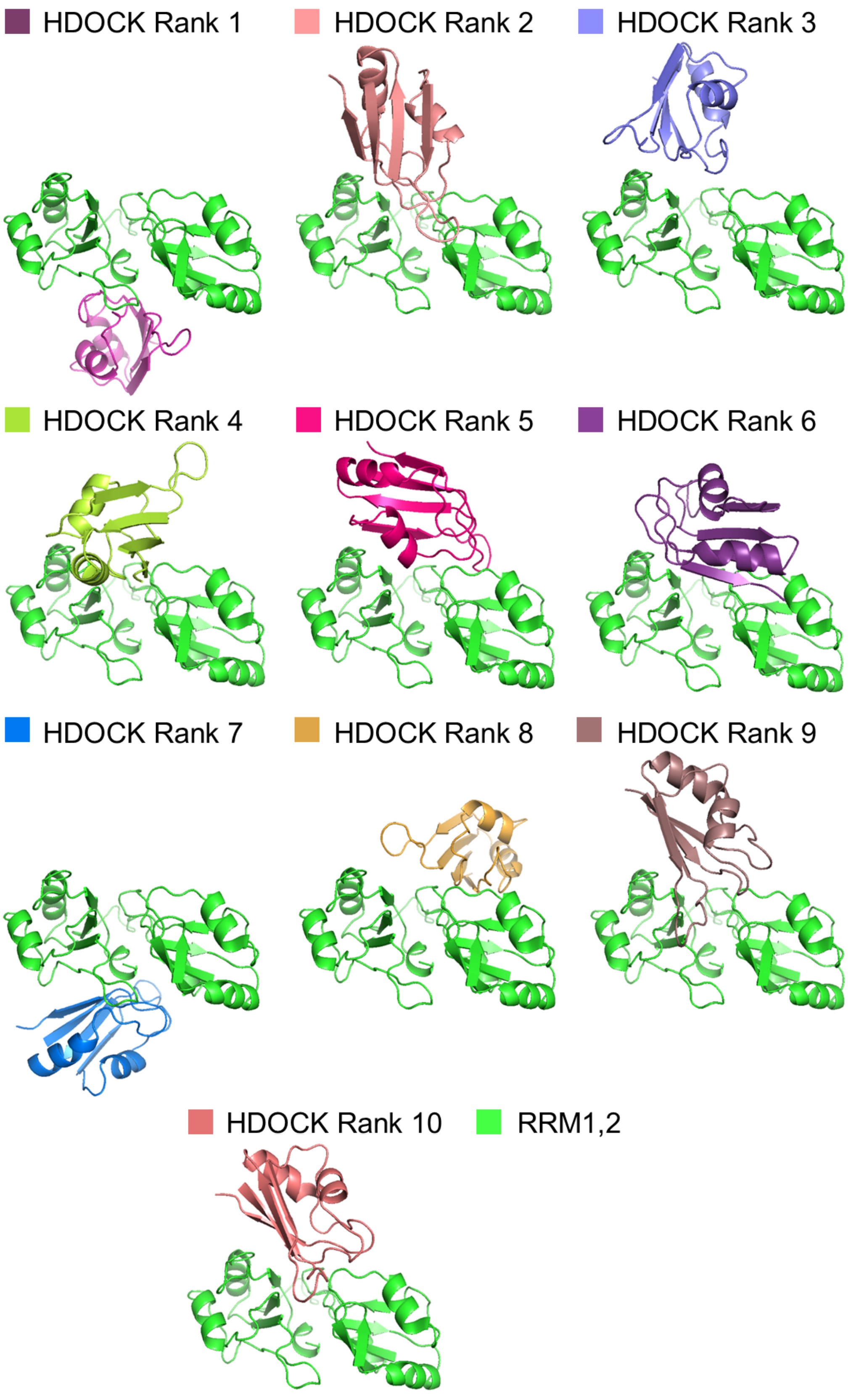


**Figure S2.** Docking poses of hnRNP H RRM3 onto hnRNP H RRMs 1 and 2.

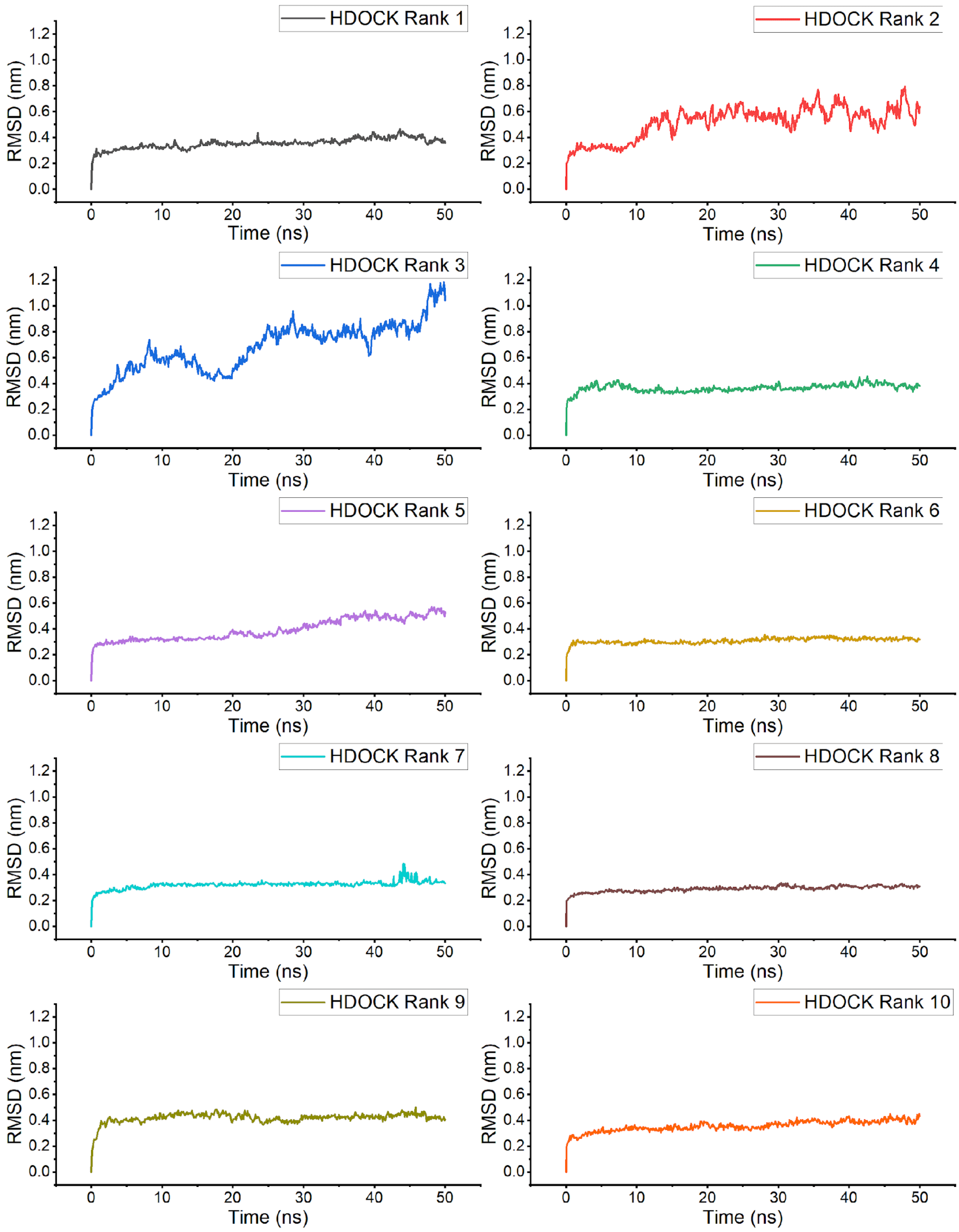


**Figure S3.** RMSD plots of the top ten docking poses of RRM3 onto RRMs 1 and 2 over 50 ns of MD simulations. RMSD calculations for each frame includes all RRMs.

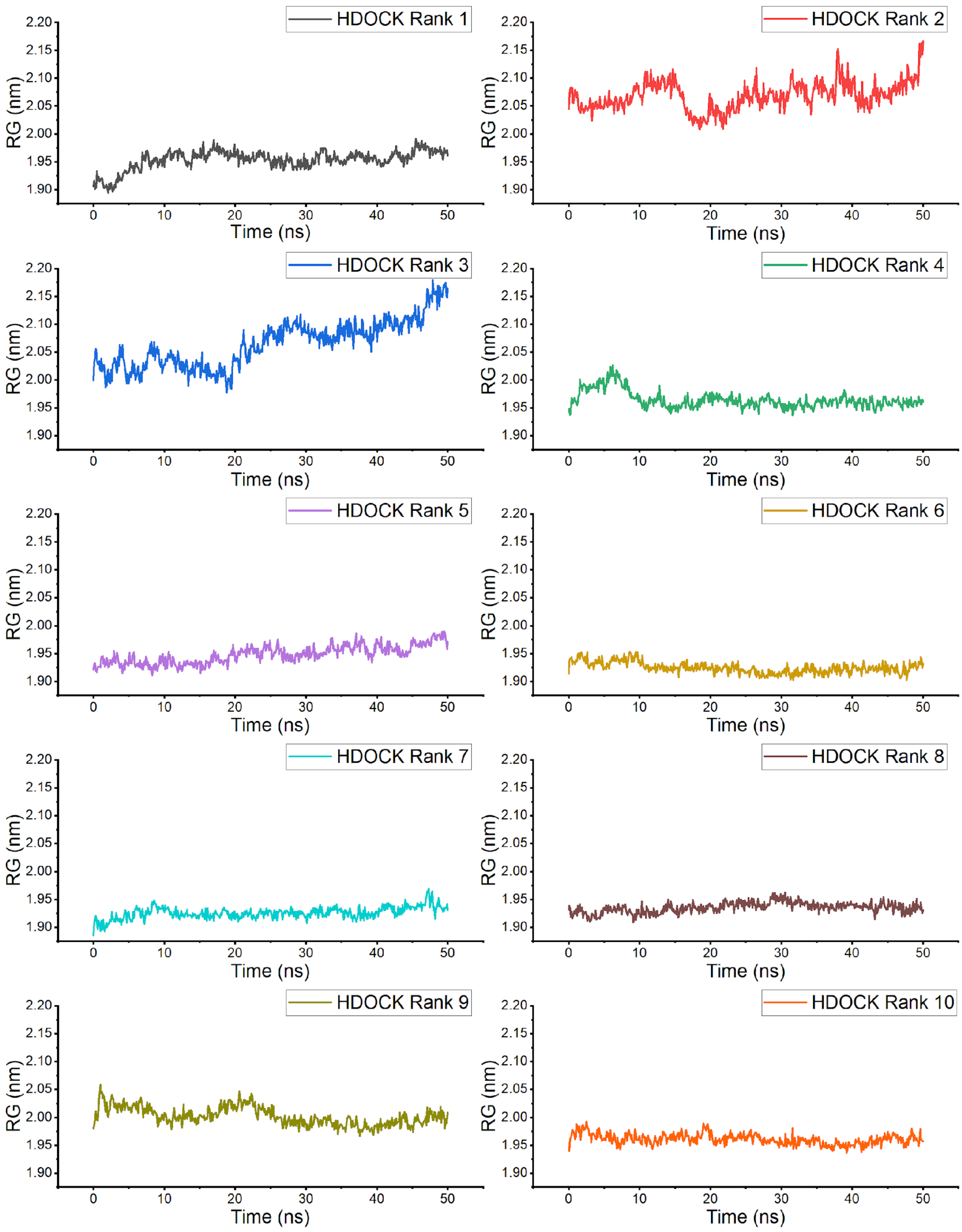


**Figure S4.** RG plots of the top ten docking poses of RRM3 onto RRMs 1 and 2 over 50 ns of MD simulations. RG calculations for each frame includes all RRMs.

Figures S3 and S4 show that the RMSD and RG for both HDOCK Ranks 2 and 3 have not stabilized within 50 ns of MD simulation. This indicates that in comparison with the other poses, which have relatively stable RMSD and RG trajectories, they are energetically unfavorable (according to the ff19SB forcefield, at least). This agrees with the MM-GBSA data presented in Table 1 and Figure 1. It can also be observed in the RMSD and RG plots for HDOCK Rank 3 that the MD simulation is deconstructing the docking pose. Figure S5 shows the 0 ns and 50 ns snapshots of HDOCK Rank 3, where it can be observed that the RRM3 moved away from RRMs 1 and 2.

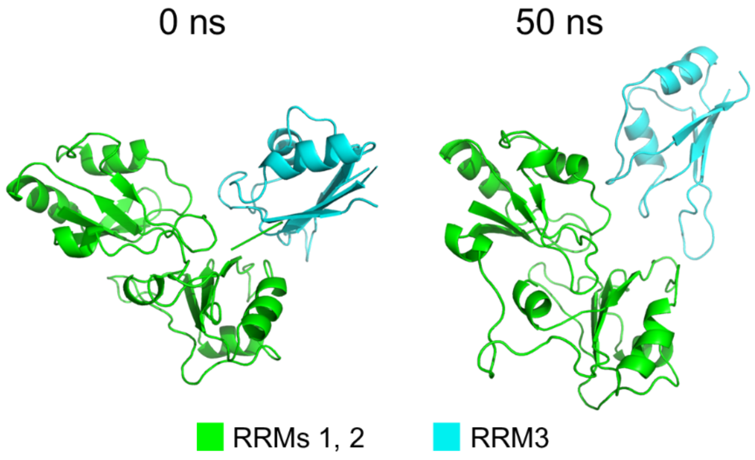


**Figure S5.** **S**napshots of HDOCK Rank 3 at 0 ns and 50 ns.

With regards to HDOCK Rank 5, from the RMSD and RG trajectories, it could be said that the structure underwent a slight reorientation but stabilized at around 38 ns. Figure S6 shows the 0 ns and 50 ns snapshots of HDOCK Rank 5, where it can be observed that the RRM3 reoriented to form more energetically favorable interactions (according to the ff19SB forcefield, at least).

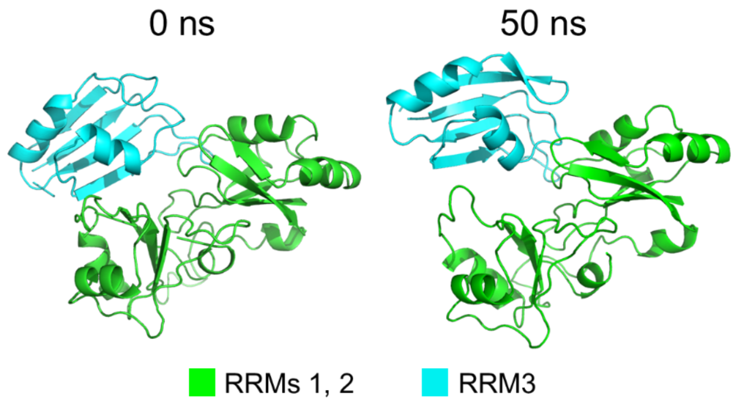


**Figure S6.** **S**napshots of HDOCK Rank 5 at 0 ns and 50 ns.

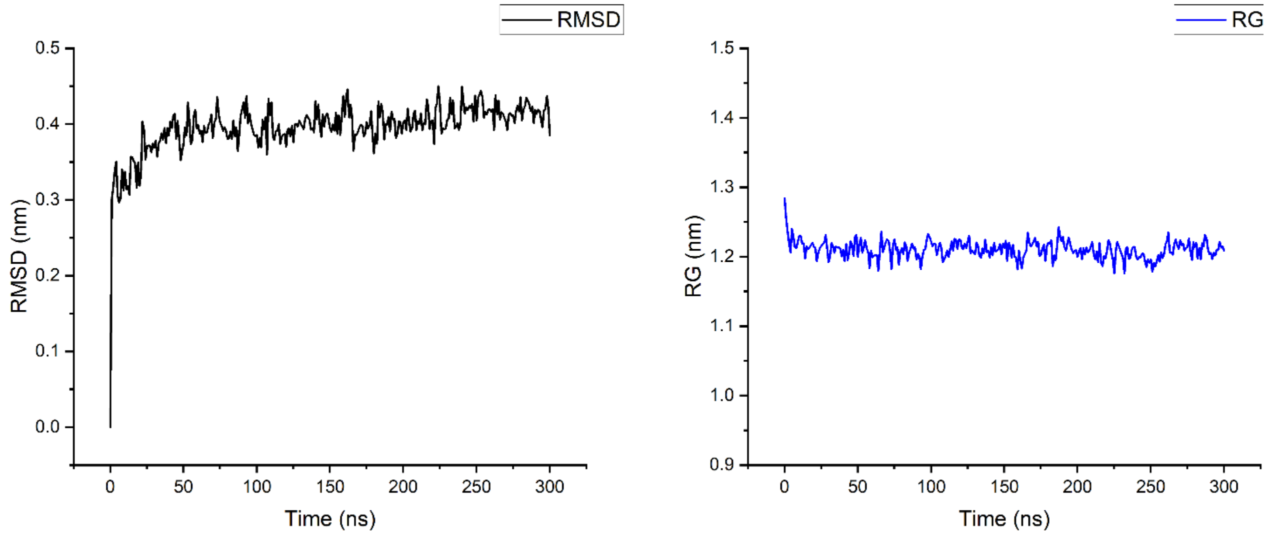


**Figure S7.** RMSD and RG plots of the MD simulation for the modelling of the C9orf72 HRE RG4. Imaginary particles incorporated for MD simulation with the Drude polarizable forcefield were not included in RMSD and RG calculations.

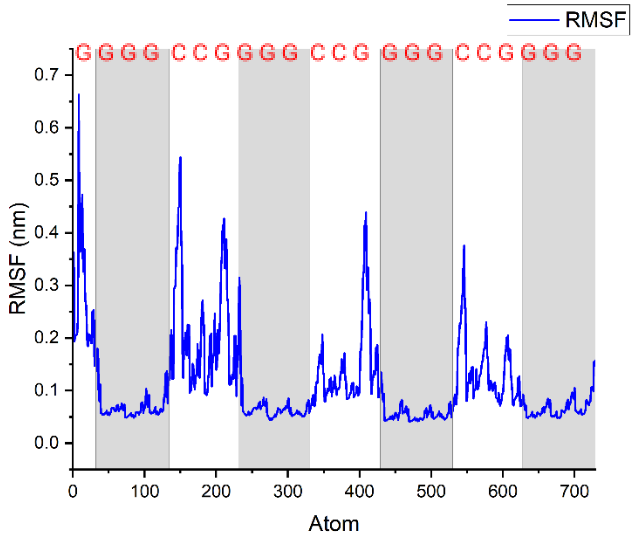


**Figure S8.** RMSF plot of the C9orf72 HRE RG4 over the last 100 ns of MD simulation for the modelling of the RG4. Highlighted regions are the guanine residues which are part of the quartets of the RG4. Imaginary particles incorporated for MD simulation with the Drude polarizable forcefield were not included in the RMSF calculations.

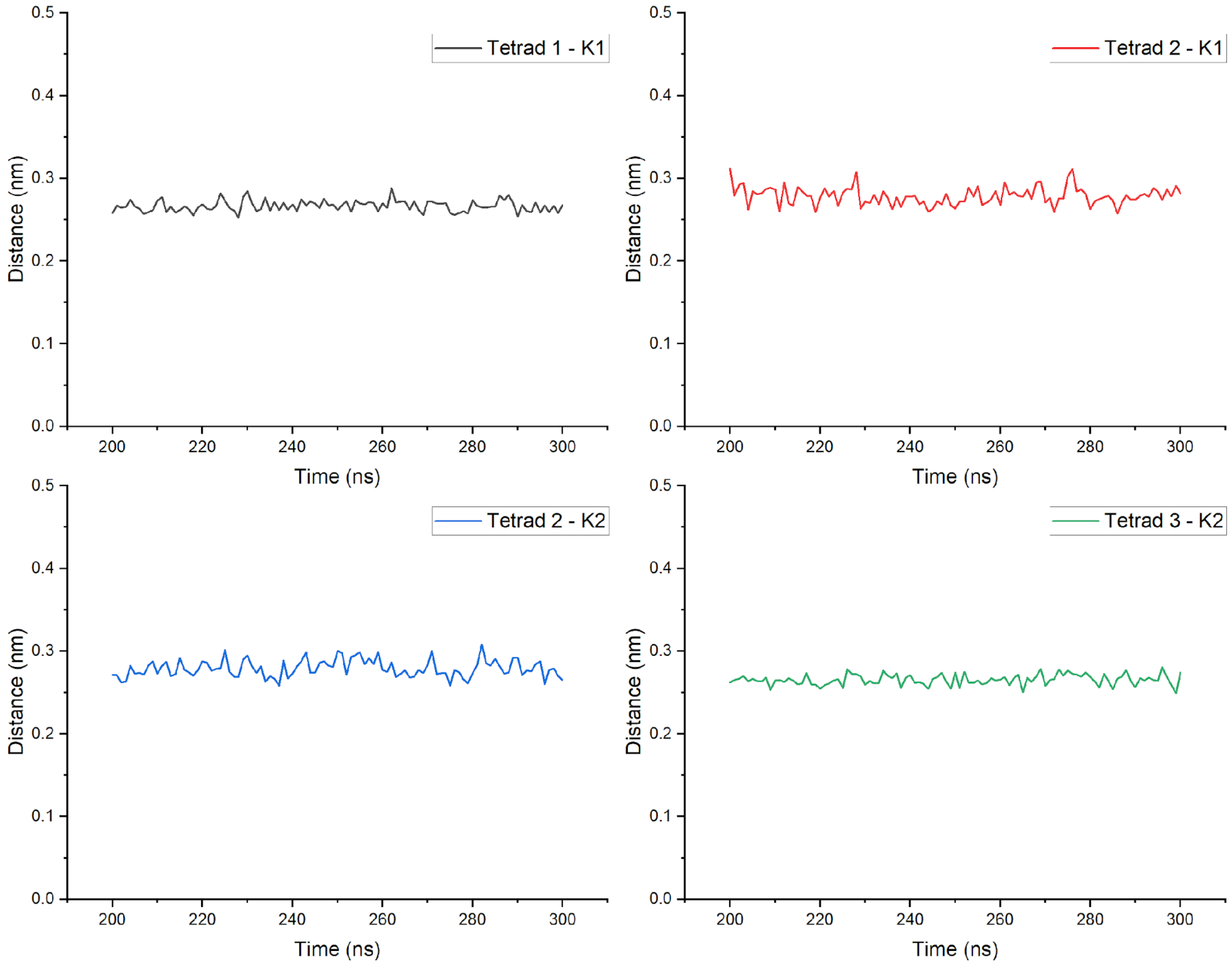


**Figure S9.** Averages of the distances of the four carbonyl oxygen atoms in the guanine bases of each tetrad to the two $K^+$ ions in the ion channel of the RG4, from 200 ns to 300 ns of the MD simulation.

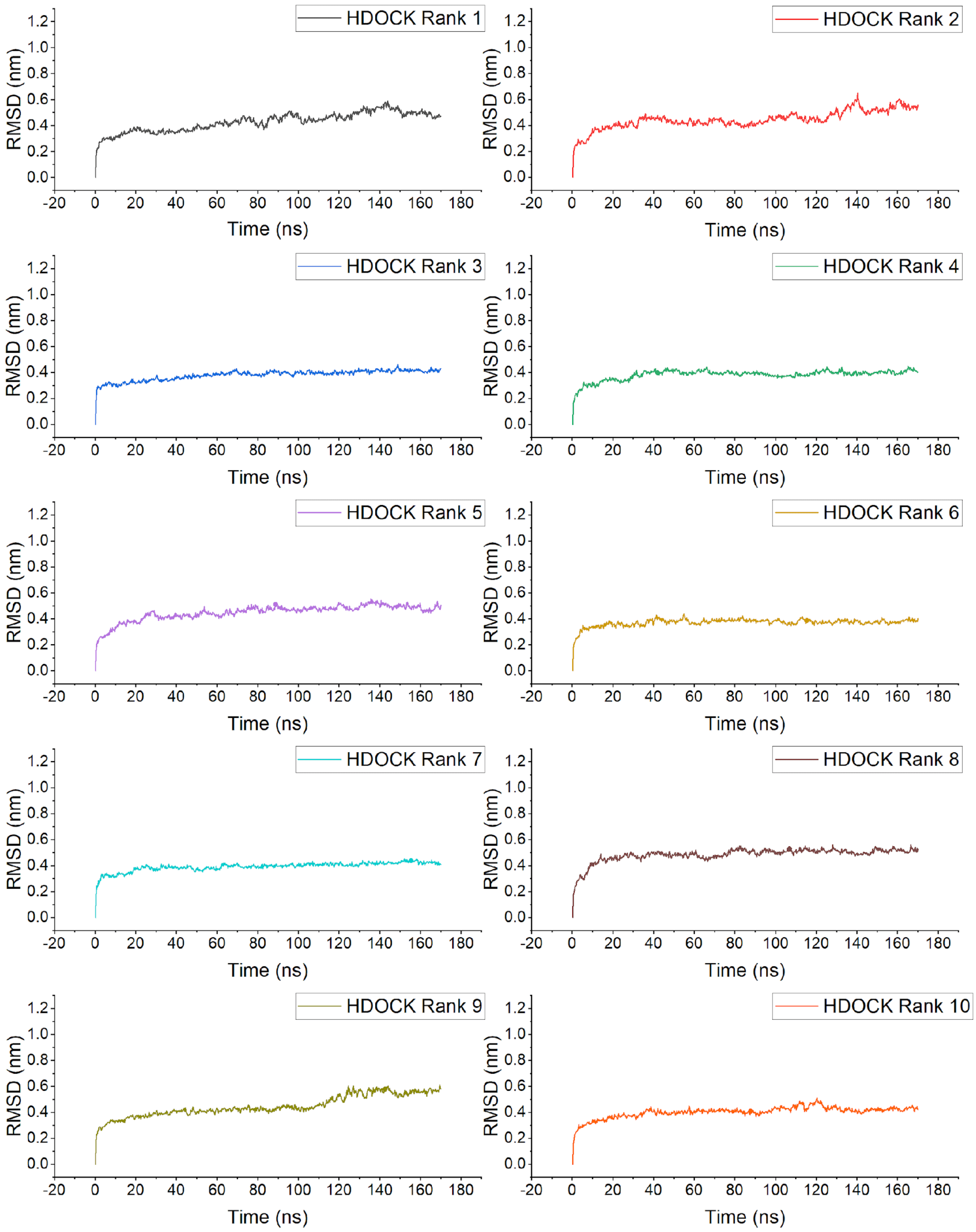


**Figure S10.** RMSD plots of the top ten docking poses of the C9orf72 HRE onto hnRNP H Model A. RMSD calculations for each frame includes both the RG4 and the hnRNP H RRMs.

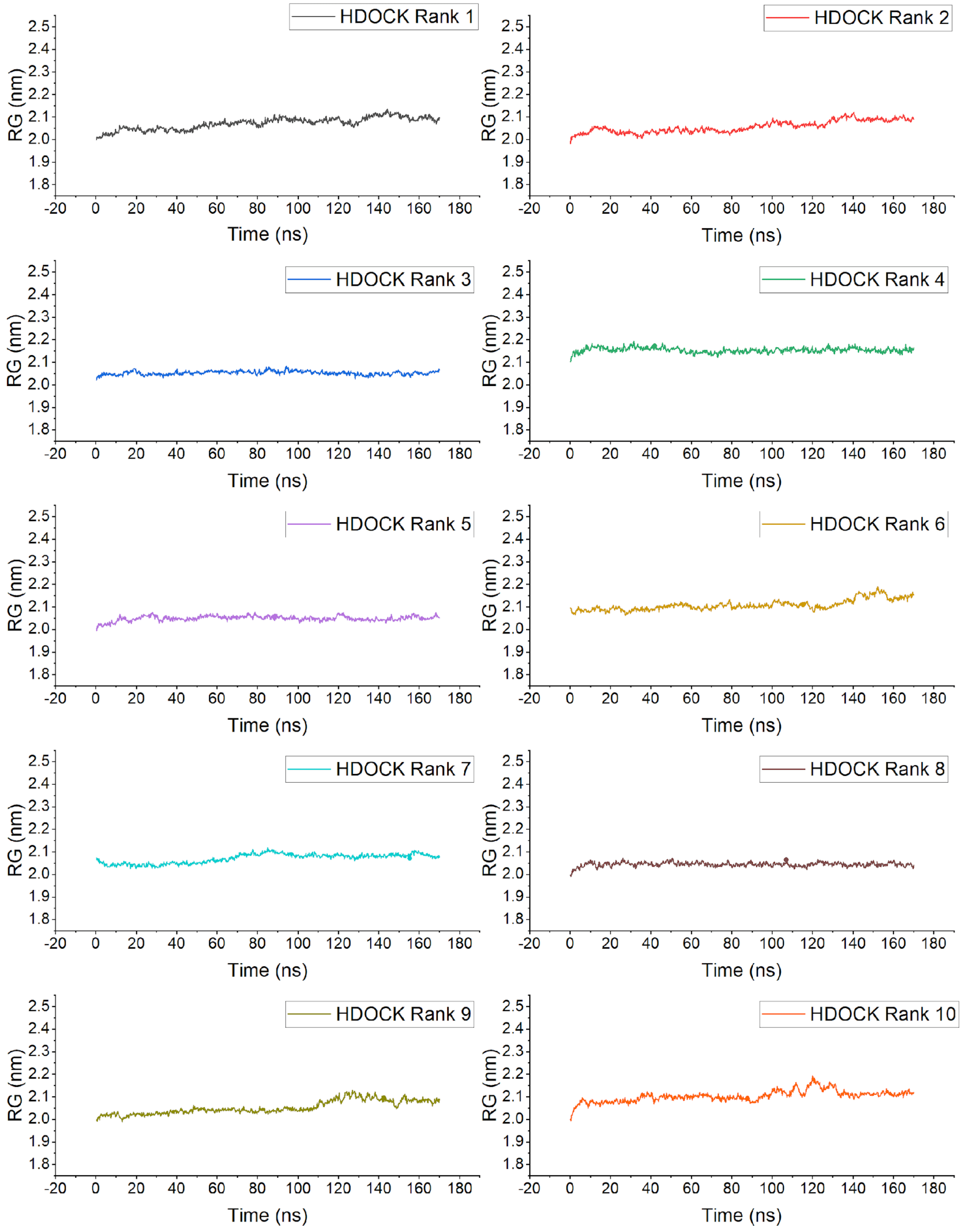


**Figure S11.** RMSD plots of the top ten docking poses of the C9orf72 HRE onto hnRNP H model A. RMSD calculations for each frame includes both the RG4 and the hnRNP H RRMs.

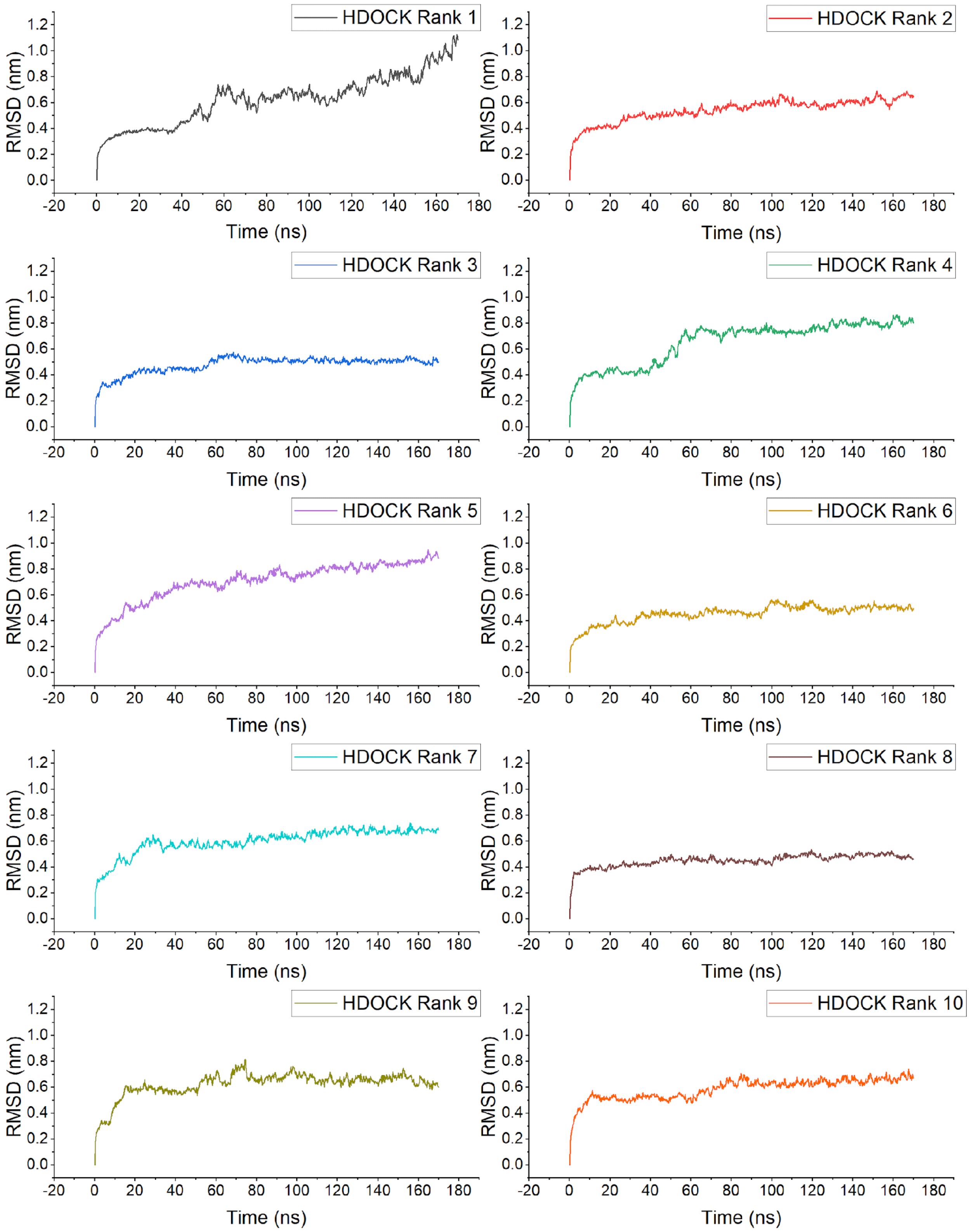


**Figure S12.** RMSD plots of the top ten docking poses of the C9orf72 HRE onto hnRNP H Model B. RMSD calculations for each frame includes both the RG4 and the hnRNP H RRMs.

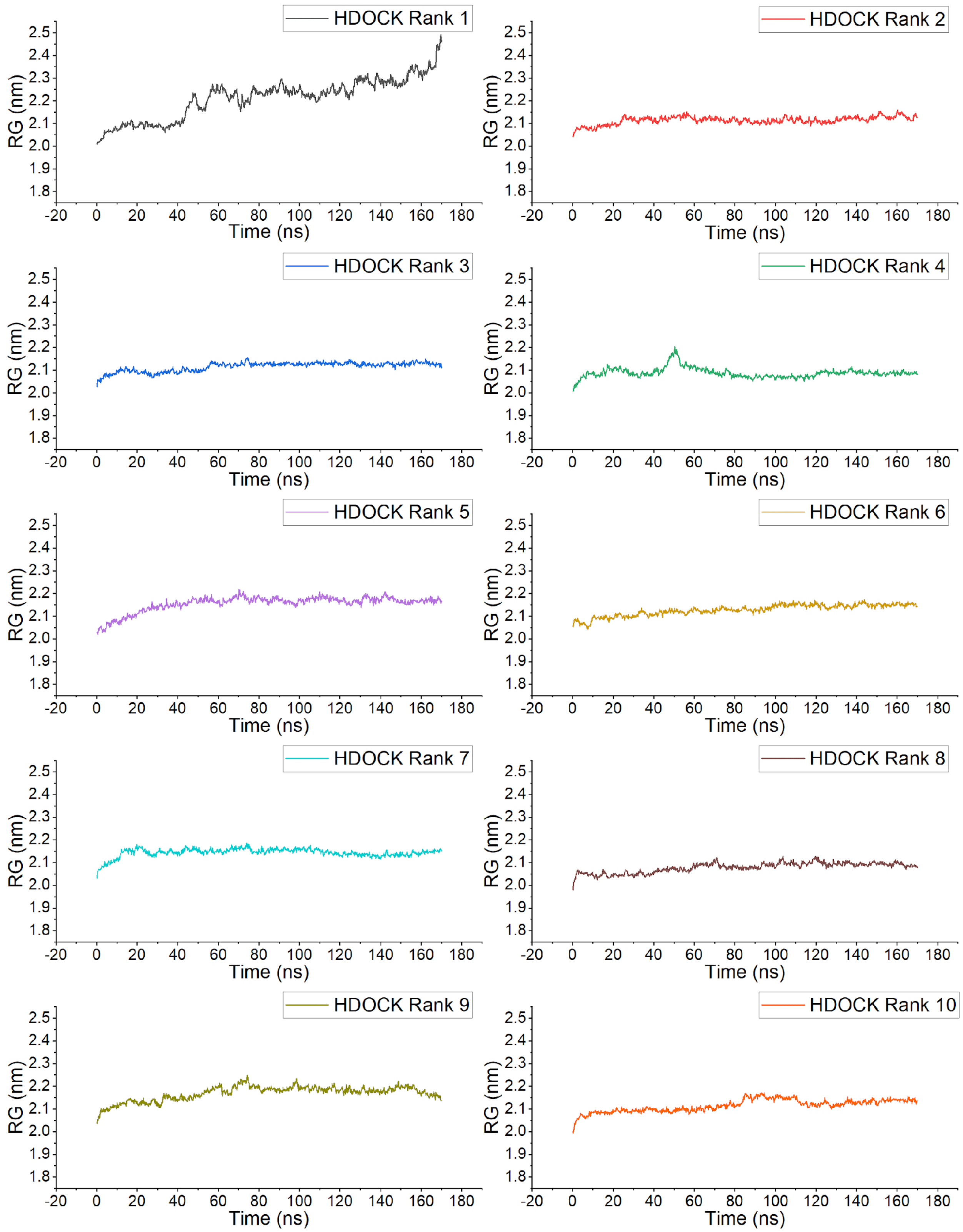


**Figure S13.** RG plots of the top ten docking poses of the C9orf72 HRE onto hnRNP H Model B. RMSD calculations for each frame includes both the RG4 and the hnRNP H RRMs.

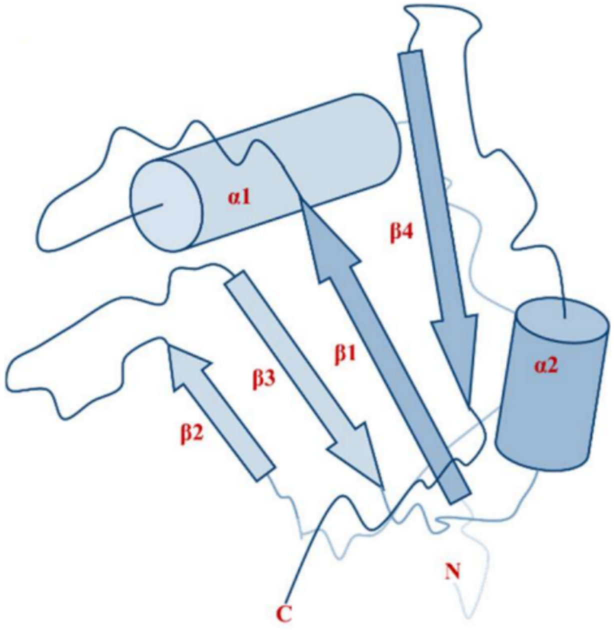


**Figure S14.** General structure of RNA recognition motifs (RRMs).

**Table S1.** Hydrogen bond analysis results for the last 30 ns of the MD simulation of the Rank 1 (global, as ranked in the MM-GBSA reranking) docking pose.

| Donor | Acceptor | Occupancy |
|---|---|---|
| THR77-Main | GUA2-Side | 53.33% |
| ARG179-Side | GUA13-Side | 36.67% |
| TYR82-Side | GUA20-Side | 23.33% |
| ARG29-Side | GUA7-Side | 23.33% |
| GUA19-Side | GLN148-Main | 21.33% |
| ARG299-Side | GUA7-Side | 13.33% |
| ARG29-Side | CYT6-Side | 12.00% |
| GLN148-Side | GUA20-Side | 5.33% |
| GLN148-Side | GUA19-Side | 5.33% |
| SER23-Side | GUA7-Side | 4.00% |
| ARG299-Side | GUA9-Side | 3.33% |
| ARG29-Side | GUA8-Side | 3.33% |
| HIS178-Side | GUA14-Side | 1.33% |
| ARG179-Side | GUA14-Side | 1.33% |
| HIS80-Side | GUA8-Side | 1.33% |
| ARG150-Side | GUA19-Side | 1.33% |
| HIS80-Side | GUA14-Side | 0.67% |
| GUA13-Side | GLY117-Main | 0.67% |
| GUA13-Side | THR152-Main | 0.67% |
| GUA7-Side | GLU26-Side | 0.67% |
| PHE120-Main | GUA14-Side | 0.67% |
| THR77-Side | GUA20-Side | 0.67% |

**Table S2.** Hydrogen bond analysis result for the last 30 ns of the MD simulation of the Rank 2 (global, as ranked in the MM-GBSA reranking) docking pose.

| Donor | Acceptor | Occupancy |
|---|---|---|
| GLN148-Side | GUA15-Side | 92.67% |
| GUA19-Side | GLN148-Side | 88.67% |
| GUA8-Side | HIS80-Side | 58.00% |
| ARG75-Side | GUA19-Side | 56.67% |
| GLN148-Side | GUA19-Side | 52.67% |
| GUA19-Side | ARG75-Main | 39.33% |
| TYR82-Side | GUA14-Side | 21.33% |
| GUA13-Side | ASP304-Side | 21.33% |
| ARG75-Side | CYT18-Side | 19.33% |
| ARG29-Side | GUA7-Side | 11.33% |
| ARG150-Side | CYT12-Side | 9.33% |
| GLN353-Side | GUA7-Side | 6.67% |
| GUA7-Side | HIS354-Side | 3.33% |
| GUA7-Side | GLN353-Side | 3.33% |
| TYR82-Side | GUA19-Side | 2.67% |
| THR77-Side | GUA14-Side | 1.33% |
| CYT12-Side | HIS178-Side | 0.67% |
| GUA14-Side | TYR82-Side | 0.67% |
| GUA19-Side | ARG75-Side | 0.67% |

**Table S3.** Hydrogen bond analysis result for the last 30 ns of the MD simulation of the Rank 3 (global, as ranked in the MM-GBSA reranking) docking pose.

| Donor | Acceptor | Occupancy |
|---|---|---|
| ARG299-Side | CYT18-Side | 74.00% |
| GUA13-Side | HIS80-Side | 46.67% |
| GUA19-Side | ARG299-Main | 40.00% |
| GUA7-Side | LYS173-Main | 25.33% |
| SER310-Side | GUA2-Side | 22.67% |
| ARG299-Side | GUA19-Side | 8.00% |
| ARG179-Side | GUA13-Side | 6.67% |
| GUA1-Side | ASP348-Side | 6.00% |
| GUA7-Side | TYR180-Side | 5.33% |
| LYS349-Side | GUA1-Side | 1.33% |
| LYS347-Side | GUA2-Side | 1.33% |
| LYS173-Side | GUA7-Side | 1.33% |
| GUA13-Side | ASN303-Side | 0.67% |
| TYR180-Side | GUA8-Side | 0.67% |
| LYS347-Side | GUA1-Side | 0.67% |
| GUA19-Side | ASP304-Side | 0.67% |

**Table S4.** Hydrogen bond analysis result for the last 30 ns of the MD simulation of the Rank 4 (global, as ranked in the MM-GBSA reranking) docking pose.

| Donor | Acceptor | Occupancy |
|---|---|---|
| ARG299-Side | CYT18-Side | 97.33% |
| ARG75-Side | GUA13-Side | 65.33% |
| ARG29-Side | GUA14-Side | 55.33% |
| TYR82-Side | GUA8-Side | 53.33% |
| ARG29-Side | GUA13-Side | 36.00% |
| LYS73-Side | GUA13-Side | 30.00% |
| GUA19-Side | GLU26-Side | 21.33% |
| ARG75-Side | GUA8-Side | 12.67% |
| ARG179-Side | GUA1-Side | 10.00% |
| CYT18-Side | GLU26-Side | 4.67% |
| HIS178-Side | GUA1-Side | 2.00% |
| HIS80-Side | GUA2-Side | 2.00% |
| GLU76-Main | GUA13-Side | 1.33% |
| ARG299-Side | GUA19-Side | 1.33% |
| ARG75-Side | GUA10-Side | 0.67% |
| TYR82-Side | GUA7-Side | 0.67% |
| GLN148-Side | GUA7-Side | 0.67% |
| GUA1-Side | GLU182-Side | 0.67% |
| ARG75-Side | GUA9-Side | 0.67% |

**Table S5.** Hydrogen bond analysis result for the last 30 ns of the MD simulation of the Rank 5 (global, as ranked in the MM-GBSA reranking) docking pose.

| Donor | Acceptor | Occupancy |
|---|---|---|
| GLN353-Side | CYT6-Side | 67.33% |
| ARG299-Side | CYT5-Side | 58.00% |
| CYT6-Side | GLN353-Main | 57.33% |
| GLN28-Side | GUA21-Side | 54.67% |
| GLN41-Main | CYT17-Side | 54.67% |
| GUA22-Side | ASP25-Main | 54.67% |
| ASN38-Side | CYT17-Side | 37.33% |
| ARG299-Side | GUA4-Side | 31.33% |
| CYT5-Side | ASP304-Side | 30.67% |
| ARG326-Side | GUA4-Side | 14.67% |
| GUA16-Side | ASP324-Side | 8.67% |
| ARG326-Side | GUA16-Side | 4.67% |
| TYR298-Side | GUA10-Side | 4.00% |
| ARG326-Side | GUA10-Side | 3.33% |
| GLN28-Side | GUA22-Side | 3.33% |
| ARG29-Side | GUA22-Side | 2.67% |
| ARG29-Side | GUA1-Side | 2.00% |
| GUA22-Side | GLN28-Side | 1.33% |
| SER32-Side | GUA1-Side | 1.33% |
| TYR298-Side | GUA4-Side | 0.67% |
| GUA21-Side | GLN28-Side | 0.67% |
| GUA1-Side | SER32-Main | 0.67% |

**Table S6.** Hydrogen bond analysis result for the last 30 ns of the MD simulation of the Rank 6 (global, as ranked in the MM-GBSA reranking) docking pose.

| Donor | Acceptor | Occupancy |
|---|---|---|
| ASN103-Side | GUA4-Side | 95.33% |
| GUA4-Side | ASN103-Side | 92.67% |
| ASN362-Side | CYT12-Side | 54.67% |
| ARG49-Side | GUA4-Side | 33.33% |
| GUA9-Side | ASN103-Side | 29.33% |
| LYS349-Side | CYT11-Side | 22.00% |
| GUA22-Side | ASP94-Side | 18.00% |
| ARG192-Side | GUA10-Side | 17.33% |
| CYT12-Side | PHE360-Main | 14.67% |
| CYT11-Side | TYR356-Side | 12.67% |
| CYT12-Side | LEU359-Main | 12.67% |
| ASN362-Side | GUA15-Side | 7.33% |
| ARG49-Side | CYT5-Side | 6.67% |
| CYT12-Side | LEU361-Main | 6.67% |
| TYR356-Side | CYT11-Side | 5.33% |
| ARG44-Side | GUA22-Side | 2.67% |
| ARG188-Side | CYT11-Side | 2.00% |
| ARG192-Side | CYT11-Side | 2.00% |
| GUA22-Side | ARG44-Side | 2.00% |
| CYT5-Side | GLU50-Main | 1.33% |
| ASN90-Side | CYT17-Side | 1.33% |
| GUA1-Side | GLN41-Side | 0.67% |
| GUA4-Side | HIS99-Main | 0.67% |
| HIS99-Side | CYT5-Side | 0.67% |

**Table S7.** Hydrogen bond analysis result for the last 30 ns of the MD simulation of the Rank 7 (global, as ranked in the MM-GBSA reranking) docking pose.

| Donor | Acceptor | Occupancy |
|---|---|---|
| ASN103-Side | GUA10-Side | 48.67% |
| SER363-Main | CYT17-Side | 46.67% |
| ASN362-Side | CYT17-Side | 45.33% |
| SER104-Side | CYT12-Side | 40.67% |
| GUA22-Side | ASP94-Side | 28.00% |
| GLN41-Side | CYT5-Side | 20.67% |
| LYS98-Side | GUA4-Side | 19.33% |
| GUA15-Side | ASN103-Side | 18.00% |
| LYS98-Side | CYT5-Side | 9.33% |
| CYT12-Side | THR107-Side | 8.67% |
| THR107-Side | CYT11-Side | 8.67% |
| CYT11-Side | ASP106-Main | 3.33% |
| GUA4-Side | HIS99-Side | 3.33% |
| TRP95-Side | GUA16-Side | 2.67% |
| CYT5-Side | GLN41-Side | 2.67% |
| SER363-Side | CYT18-Side | 2.67% |
| HIS99-Side | CYT11-Side | 2.67% |
| LYS98-Side | CYT6-Side | 2.00% |
| CYT17-Side | LEU361-Main | 0.67% |
| SER363-Side | CYT17-Side | 0.67% |
| CYT12-Side | SER104-Side | 0.67% |
| GUA10-Side | HIS99-Side | 0.67% |

**Table S8.** Hydrogen bond analysis result for the last 30 ns of the MD simulation of the Rank 8 (global, as ranked in the MM-GBSA reranking) docking pose.

| Donor | Acceptor | Occupancy |
|---|---|---|
| GUA7-Side | ASP304-Side | 90.00% |
| GUA19-Side | THR77-Side | 86.67% |
| GUA13-Side | LYS72-Main | 84.00% |
| GUA13-Side | GLU76-Side | 78.00% |
| PHE120-Main | GUA20-Side | 75.33% |
| ARG150-Side | GUA19-Side | 60.00% |
| GUA19-Side | HIS80-Main | 57.33% |
| GUA1-Side | GLU125-Side | 14.00% |
| ARG299-Side | GUA8-Side | 10.00% |
| GUA19-Side | ARG150-Side | 8.00% |
| CYT18-Side | GLY79-Main | 4.67% |
| CYT18-Side | HIS80-Side | 4.67% |
| TYR82-Side | GUA13-Side | 3.33% |
| GUA13-Side | ARG75-Main | 2.67% |
| HIS80-Side | CYT18-Side | 0.67% |
| GLN148-Side | GUA8-Side | 0.67% |
| THR77-Side | GUA14-Side | 0.67% |
| HIS80-Side | GUA19-Side | 0.67% |
| GUA19-Side | GLY79-Main | 0.67% |

**Table S9.** Hydrogen bond analysis result for the last 30 ns of the MD simulation of the Rank 9 (global, as ranked in the MM-GBSA reranking) docking pose.

| Donor | Acceptor | Occupancy |
|---|---|---|
| ARG29-Side | CYT18-Side | 88.67% |
| HIS80-Side | GUA2-Side | 61.33% |
| GUA13-Side | ARG29-Main | 58.00% |
| ARG29-Side | GUA19-Side | 55.33% |
| ARG150-Side | GUA2-Side | 53.33% |
| GUA2-Side | TYR82-Side | 46.00% |
| ARG29-Side | GUA14-Side | 30.67% |
| TYR82-Side | GUA2-Side | 26.67% |
| GUA19-Side | ASP25-Side | 19.33% |
| SER23-Side | GUA19-Side | 12.00% |
| GUA13-Side | PHE30-Main | 12.00% |
| ARG75-Side | GUA7-Side | 6.67% |
| SER21-Side | GUA20-Side | 4.00% |
| GUA19-Side | GLU26-Side | 2.00% |
| GLN148-Side | GUA7-Side | 0.67% |

**Table S10.** Hydrogen bond analysis result for the last 30 ns of the MD simulation of the Rank 10 (global, as ranked in the MM-GBSA reranking) docking pose.

| Donor | Acceptor | Occupancy |
|---|---|---|
| LYS124-Main | GUA13-Side | 84.00% |
| ARG75-Side | GUA19-Side | 47.33% |
| LYS124-Side | GUA13-Side | 44.67% |
| GUA13-Side | CYS122-Main | 38.67% |
| GUA13-Side | GLY149-Main | 32.00% |
| GUA7-Side | GLU126-Side | 18.00% |
| ARG299-Side | GUA15-Side | 17.33% |
| GLN129-Side | GUA7-Side | 17.33% |
| GUA19-Side | ARG75-Side | 12.67% |
| GLN148-Side | GUA19-Side | 12.00% |
| GLY177-Main | GUA7-Side | 10.67% |
| SER123-Side | GUA13-Side | 7.33% |
| GLN148-Side | GUA14-Side | 3.33% |
| ARG299-Side | GUA16-Side | 2.67% |
| GUA7-Side | GLN129-Side | 2.00% |
| SER123-Side | GUA8-Side | 1.33% |
| ARG75-Side | GUA20-Side | 1.33% |
| GLN148-Side | GUA20-Side | 1.33% |
| ARG299-Side | GUA14-Side | 0.67% |
| GUA19-Side | TYR298-Main | 0.67% |
| GUA13-Side | GLY121-Main | 0.67% |

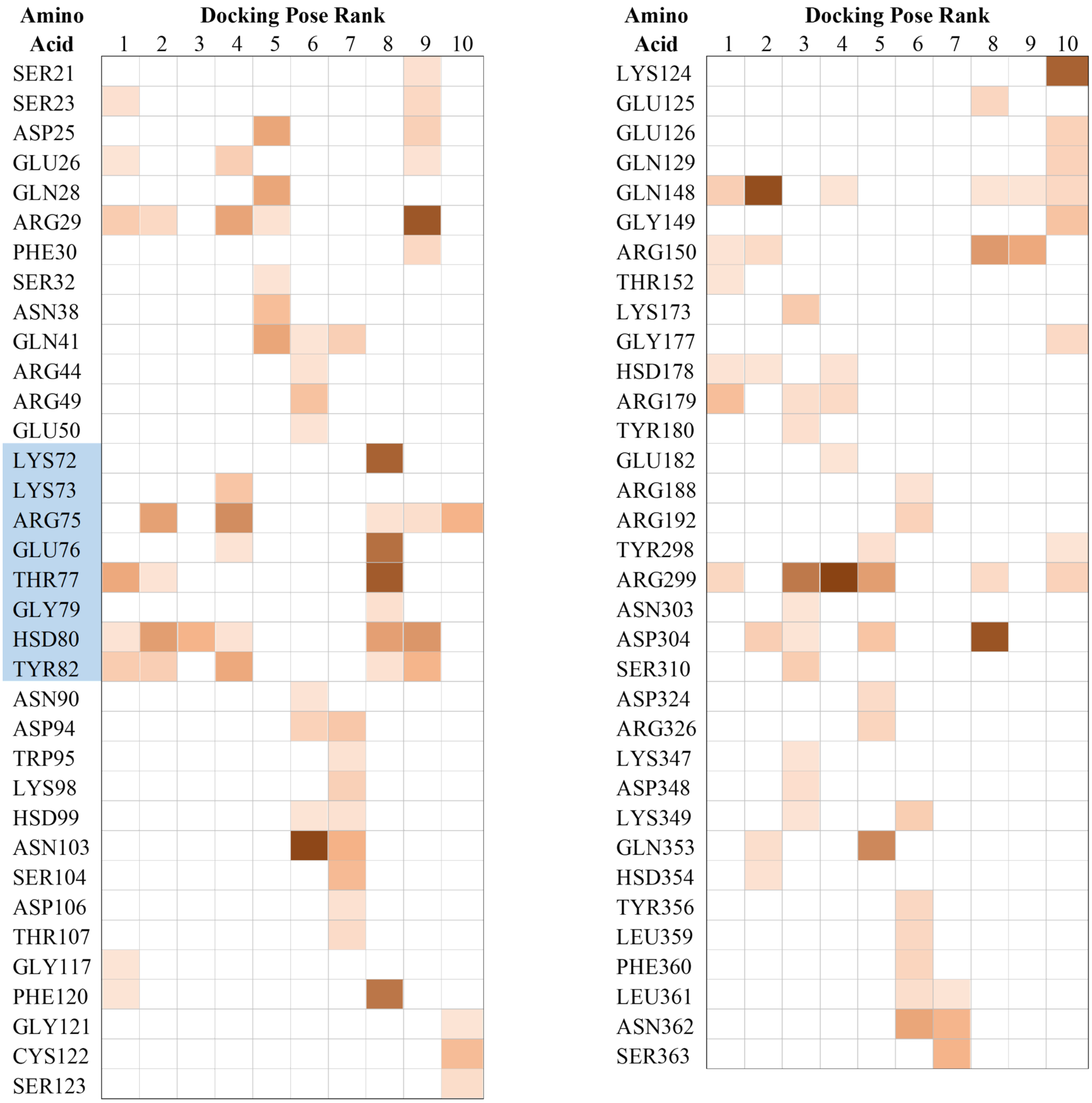


**Figure S15.** Graphic summary of the data presented in Tables S1 to S10. Intensity of the cell highlight represents the occupancy of the most prominent hydrogen bond that the amino acid residues of the hnRNP H make with the RG4 (more intense means higher occupancy). Amino acid residues highlighted in blue are proposed to be a region of interest in studying the interaction of hnRNP H with the C9orf72 HRE RG4.
